# Transgenerational Regulation of Sexual Attractiveness in *C. elegans* Nematodes

**DOI:** 10.1101/2020.11.18.389387

**Authors:** Itai Antoine Toker, Itamar Lev, Yael Mor, Yael Gurevich, Doron Fisher, Leah Houri-Zeevi, Olga Antonova, Lilach Hadany, Shai Shaham, Oded Rechavi

## Abstract

It is unknown whether transient transgenerational epigenetic responses to environmental challenges affect the process of evolution, which typically unfolds over many generations. Here we show that in *C. elegans*, inherited small RNAs shape the hard-wired genome and control genetic variation by regulating the decision of whether to self-fertilize or outcross. We found that under stressful temperatures younger hermaphrodites secrete a male-attracting pheromone. Attractiveness transmits transgenerationally to unstressed progeny via heritable small RNAs and the Argonaute Heritable-RNAi-Deficient-1. We identified an endogenous small interfering RNA pathway, enriched in endo-siRNAs which target sperm genes, that transgenerationally regulates sexual attraction, male prevalence, and outcrossing rates. Multigenerational mating competitions and mathematical simulations revealed that over generations, animals that inherit attractiveness mate more, and their alleles spread in the population. We propose that sperm serves as a “stress sensor” which, via small RNA inheritance, can promote outcrossing in challenging environments, when increasing genetic variation is advantageous.

## Introduction

Hermaphrodites of the androdioecious nematode species *Caenorhabditis elegans* can either self-reproduce or outcross. Outcrossing comes at great cost, as it requires finding a partner, exposes the animals to physical damage, increases predation risk, and, importantly, dilutes each parent’s genetic contribution(*1*). However, theory predicts that increased tendency for outcrossing under stress would evolve in many cases, and could facilitate adaptation(*2*, *3*). Accordingly, it was shown that *C. elegans* outcross more when exposed to stressful conditions(*4*, *5*), and that the genetic variability gained following outcrossing is essential for adaptation to changing environments(*6*). Recent studies revealed that when aging *C. elegans* hermaphrodites run out of self-made sperm and lose the ability to self-reproduce, they secrete a volatile sex pheromone that elevates the chances of mating with males(*7*, *8*). The secretion of this pheromone is triggered when there is no longer contact between the depleted sperm and the oocytes in the gonads(*7*). The exact chemical nature of the pheromone is unknown, however it was shown that males detect it using the G protein-coupled receptor SRD-1 in the bilateral AWA sensory neurons(*8*).

Inheritance of experiences has been a controversial topic for centuries(*9*, *10*). Nevertheless, when *C. elegans* are stressed they can generate physiological responses that carry on transgenerationally(*11–18*). Some of the effects were shown to transmit independently of DNA changes by heritable small RNAs which can be re-synthesized and amplified in the progeny by RNA-dependent RNA polymerases(*11*, *19*). Dedicated machinery regulates the transmission and maintenance of these effects(*20–28*), and disruption of small RNA inheritance results in damages to the germ cells that accumulate over generations and can lead to full sterility(*20*, *23*, *24*, *29*, *30*). The germline, therefore, both serves as a rheostat that controls the secretion of male-attracting pheromones, and reflects the effects of environmental responses on the pools of heritable small RNAs. Hence, we hypothesized that the secretion of the male-attracting pheromone might be induced by environmental stress and memorized by heritable small RNAs.

## Results

### Environmental stress induces transgenerational premature attraction via heritable small RNAs

To test our hypothesis, we first performed chemotaxis experiments to measure male attraction to odors extracted from the growth medium of one-day-old hermaphrodites exposed to different environmental stresses (Fig. 1A). Under standard growth conditions, wild-type hermaphrodites do not secrete the male-attracting pheromone at this young age. As a positive control we used odors extracted from *fog-2(q71)* adult hermaphrodites that do not produce sperm and are therefore constitutively attractive(*7*, *8*, *31*).

**Fig.1.**
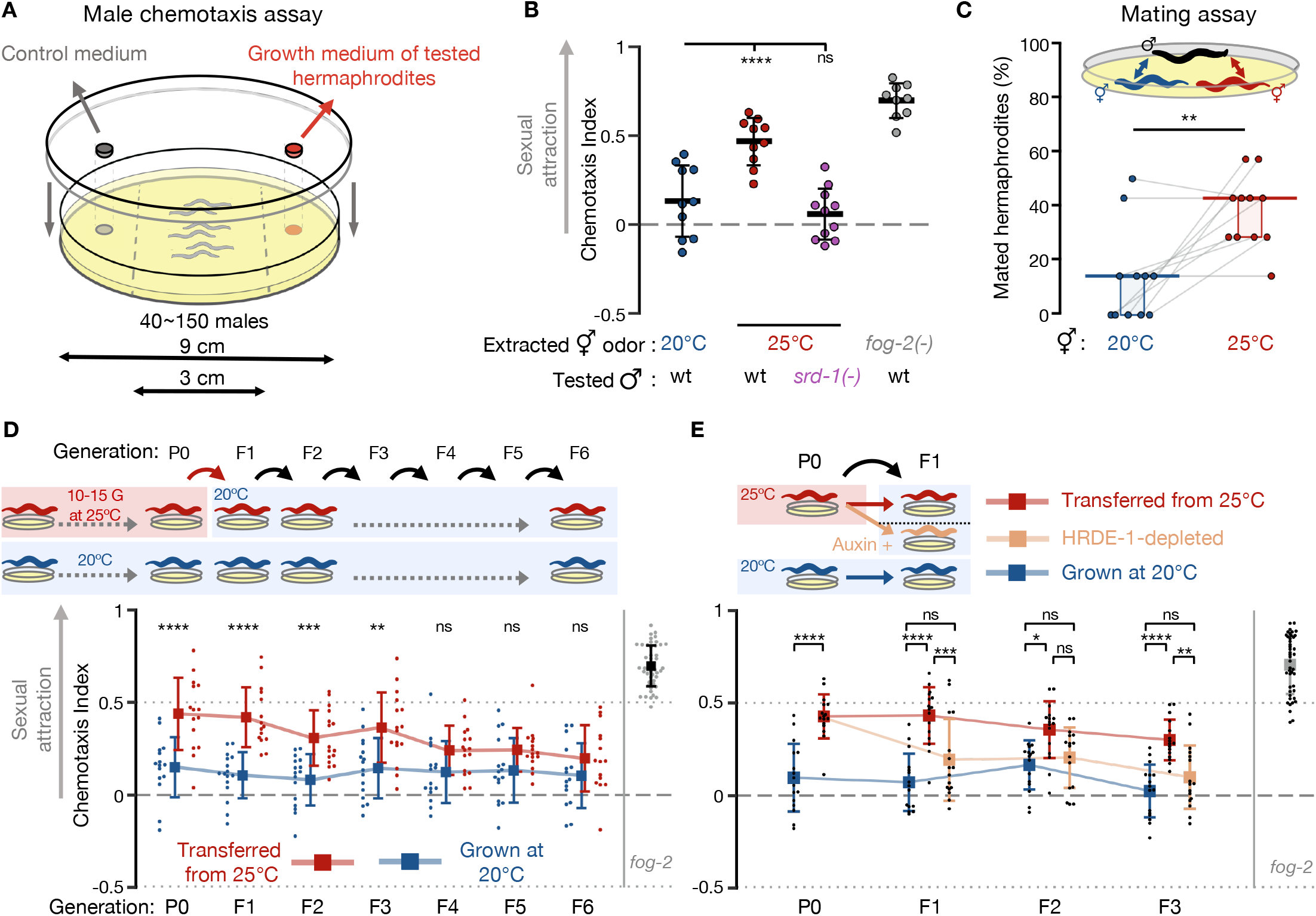
Environmental stress induces transgenerational premature attraction via HRDE-1-dependent heritable small RNAs. (**A**) Scheme of the male volatile chemotaxis assay. (**B**) Male chemotaxis experiments. Growth temperature of hermaphrodites used for odor extraction and males used for chemotaxis appear below the panel. Each dot represents one biological replicate (chemotaxis plate) with 37-152 males. Bars: Mean ± sd, results from 3 independent experiments. One-way ANOVA, Dunnett’s correction for multiple comparisons to 20°C. (**C**) Proportions (%) of hermaphrodites mated during mating choice experiments. Young hermaphrodites previously cultivated at the indicated temperatures interacted with wild-type males for exactly 1 hour. Each grey line represents one biological replicate, results were collected over three independent experiments. Horizontal bar: Median. Boxes: Interquartile range (IQR). Two-tailed Wilcoxon matched-pairs signed rank test. See negative (non-attracting) and positive (*fog-2* mutants) control groups in Fig. S2. (**D**) Male chemotaxis experiments with odors extracted from hermaphrodites grown at 25°C for 10-15 generations (P0) and transferred back to 20°C for 6 generations (red). Continuously 20°C-grown hermaphrodites were used as control (blue). Each dot represents one biological replicate (chemotaxis plate) with 38-144 males. Bars: mean ± sd, results from 4 independent experiments. Two-way ANOVA, Sidak’s correction for multiple comparisons for every generation. (**E**) Odors were extracted from 25°C-grown *aid::hrde-1;sun-1p::tir-1* worms cultivated with (beige) or without auxin (red). Experiment was performed with additional control groups (see Fig. S4A) accounted for in the presented statistical analysis. Each dot represents one biological replicate (chemotaxis plate) with 38-138 males. Bars: mean ± sd, results from 4 independent experiments. Two-way ANOVA, Tukey’s correction for multiple comparisons. (**B-E)** \*\*\*\**P* < 10^−4^, \*\*\**P* < 0.001, \*\**P* < 0.01, \**P* < 0.05, ns *P* > 0.05.

We examined a few stressful conditions: short-term incubation at 25°C, short-term starvation at L1 larval stage and long-term starvation in the dormant *dauer* stage. None of these conditions induced premature attractiveness (Fig. S1A). Previous studies have shown that temperature-induced gene expression changes accumulate over generations(*14*, *32*). When we continuously cultivated the animals for 10-15 generations at 25°C, which is mildly stressful but within the standard temperature range for lab cultures, we discovered that the adult hermaphrodites were prematurely attractive (Fig. 1B). Attractiveness seemed to accumulate over generations among hermaphrodites grown at 25°C for 1, 4, 8 & 12 generations (Fig. S1B). Attraction required the activity of the pheromone receptor SRD-1 in the males (Fig. 1B), indicating that this premature attractiveness is detected by the same sensory pathway that mediates attraction towards aged or spermless hermaphrodites(*8*). To examine if premature attractiveness translates also to an increase in mating, we performed mating choice assays in which males carrying fluorescently labeled sperm had to choose between synchronized control (grown at 20°C) and prematurely attractive hermaphrodites (grown at 25°C). In the physical presence of both mating partners, the males copulated more with the hermaphrodites grown at 25°C (14% vs 43% mated hermaphrodites, Fig. 1C, *P*=0.008, and Fig. S2 for controls). The increase in insemination rates was *srd-1*-independent under these co-culture conditions (Fig. S2), suggesting that distinct locally-acting cues, in addition to the secreted volatile odor, could further stimulate the males or the stressed hermaphrodites to mate(*33–37*).

Strikingly, we found that premature attractiveness was maintained transgenerationally for three generations in descendants transferred to standard growth conditions (20°C) (Fig. 1D), before it petered out. How is premature attractiveness inherited across generations? We hypothesized that this process could be mediated by changes in heritable small RNAs, since germline small RNAs are modulated in response to high temperatures(*13*, *32*) and regulate key germline functions(*38–41*). The inheritance of many germline endogenous small interfering RNAs (endo-siRNAs)(*13*, *19*, *20*), and the transgenerational inheritance of different small RNA-mediated phenotypes(*12*, *42–45*), require the Argonaute HRDE-1 (**H**eritable **R**NAi **De**ficient-1). Because *hrde-1* mutants are sterile when cultivated at 25°C(*20*, *23*), we used CRISPR-Cas9 to engineer strains permitting conditional HRDE-1 depletion via the auxin-inducible degradation (AID) system(*46*) (see Methods and Fig. S3). By depleting HRDE-1 in descendants grown at 20°C we revealed that the heat (25°C)-triggered inheritance of premature attractiveness depends on HRDE-1 and therefore on small RNA inheritance (Fig. 1E and see Fig. S4A for additional controls). The Histone-3-Lysine-9 histone-methyltransferase SET-25 has been previously shown to be involved in certain transgenerational temperature-induced effects(*14*), but our results indicate that it is not required for the inheritance of premature attractiveness (Fig. S4B). Together, these results demonstrate that exposure to environmental stress induces enhanced sexual attractiveness in young hermaphrodites, which is transmitted transgenerationally via the small RNA inheritance machinery.

### MEG-3/4-dependent small RNAs regulate heritable sexual attraction

To dissect the small RNA pathways involved in heritable premature attractiveness, we assayed males for chemotaxis towards odorants extracted from one-day-old hermaphrodites carrying lesions in genes required in different small RNA pathways (Fig. 2A and Table S1). We tested 15 small RNA mutant strains and discovered that *meg-3(tm4259)*;*meg-4(ax2026)* double mutants are prematurely attractive to males in a SRD-1-dependent manner, and moreover, that they transmit the premature attractiveness transgenerationally to their wild-type descendants (Fig. 2B,C). Three other strains, *prg-1(n4357), dcr-1(mg375)* & *alg-5(tm1163)*, were found to be prematurely attractive, however they did not transmit the attractiveness transgenerationally (Fig. 2A and Fig. S5). In addition to secreting the attractive odors, hermaphrodites from these four strains mated more with males in mating choice assays (Fig. 2D and Fig. S6).

**Fig.2.**
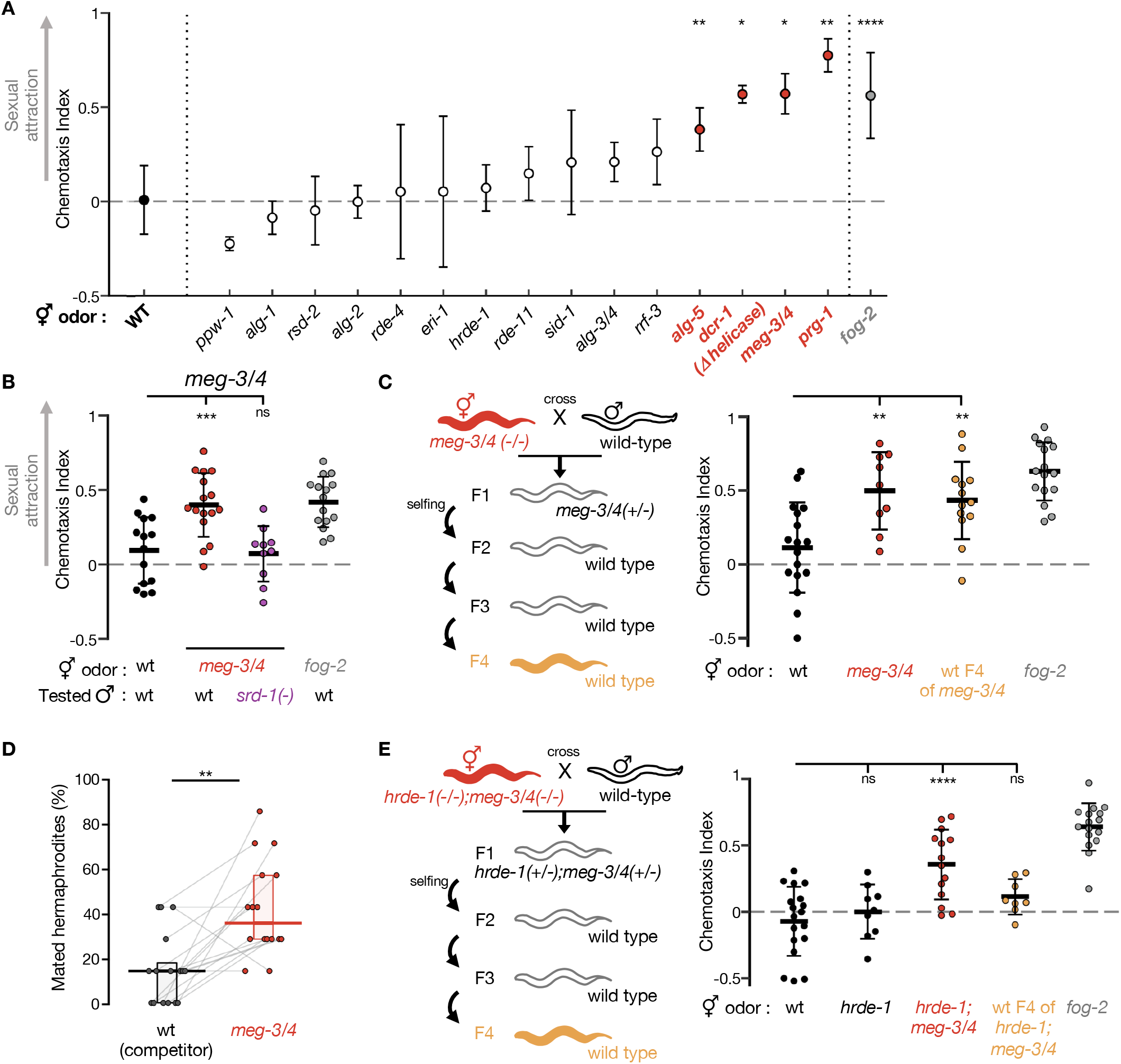
MEG-3/4-dependent small RNAs regulate heritable sexual attraction. **(A)** Premature attractiveness screening of mutants impaired in genes acting in different small RNA pathways. Odors were extracted from mutated (x-axis) young hermaphrodites, and tested for the attraction of wild-type males. Mean ± sd. Kruskal-Wallis test with Dunn’s correction for multiple comparisons to wild-type. Full information appears in Table S1. (**B**) Male chemotaxis experiments. Genotypes of hermaphrodites used for odor extraction and of tested males appear below panel. (**C**) Male chemotaxis experiments testing for odor extracted from young wild-type descendants of *meg-3/4* double mutants. (**D**) Proportions (%) of mated hermaphrodites during 1hour-mating choice experiments, genotype indicated below panels. Grey lines represent biological replicates, results collected over 4 independent experiments. Horizontal bar: Median. Boxes: IQR. Two-tailed Wilcoxon matched-pairs signed rank test. (**E**) Male chemotaxis experiments testing for odor extracted from wild-type descendants of *hrde-1;meg-3/4* triple mutants. (**C,E)** Tested odors are color-coded according to schemes, and all hermaphrodites contained an integrated single-copy *mex-5p::gfp* transgene. (**B,C,E**) Each dot represents one biological replicate (chemotaxis plate) with 38-190 males. Bars: mean ± sd. One-way ANOVA, Dunnett’s correction for multiple comparisons to wild-type. All conditions were tested across at least 3 independent experiments. \*\*\*\**P* < 10^−4^, \*\*\**P* < 0.001, \*\**P* < 0.01, \**P* < 0.05, ns *P* > 0.05.

Similarly to the transgenerational effect of long-term growth at 25°C, we found that the inheritance of premature attractiveness from *meg-3/4* mutants to wild-type progeny depended on the Argonaute HRDE-1 (Fig. 2E). The heritable attractiveness of *meg-3/4* was particularly potent and lasted for six generations (Fig. S7). This result resonates with previous findings showing that *meg-3/4* mutants exhibit a transgenerational disruption in the expression patterns of small RNAs pools in the germline(*43*, *47*, *48*). These misregulated heritable small RNAs lead to extraordinarily strong inheritance of RNAi defects that can be detected in their wild-type descendants even after >10 generations(*43*, *47*).

Interestingly, all the RNA processing proteins that we found to affect premature attractiveness are located in the germ granules(*49–53*), which are cytoplasmic condensates made of RNA and proteins found in the germline of a great number of organisms. *meg-3/4* double mutants display severe defects in germ granules(*53*). Moreover, and in accordance with a previous study(*54*), we found that exposure to heat reduced the size of the P granules in the adult germline (Fig. S8). As was previously shown(*43*, *47*, *48*), transiently disrupting the germ granules’ stability can have major heritable consequences on germline gene expression, as it dramatically alters the pool of heritable small RNAs (See more below). Thus, the small RNA-mediated heritable attractiveness of worms subjected to 25°C may derive from the effect of elevated temperatures on the stability of germ granules and on the processing of germline small RNAs therein.

### Premature attractiveness is associated with misregulation of sperm genes and sperm defects

How do heritable small RNAs control attraction? Numerous endo-siRNAs, emanating from throughout the genome, are misregulated in worms grown at 25°C for multiple generations, in *meg-3/4* mutants, and in wild-type descendants of *meg-3/4* mutants(*13*, *32*, *43*, *47*, *48*). Our analyses revealed that the target genes of these endo-siRNAs, and the mRNAs differentially-expressed under these conditions, are enriched for sperm-expressed genes (Fig. 3A and Fig. S9A). A subset of 320 protein-coding genes were targeted by small RNAs that transgenerationally accumulate in worms cultivated at 25°C, and these genes were also enriched for sperm-expressed genes (Fig S9B). Wild-type animals grown at 25°C for 10 generations were previously shown to have sperm defects(*32*, *38*, *55*). We found that both *meg-3/4* mutants, and their wild-type descendants, exhibit a reduction in selfing-derived brood size (−22%, −13%, Fig. S10A). In mating rescue experiments (Fig. 3B), the brood size of *meg-3/4* mutants robustly increased following mating with wild-type males (2.2x-fold, *P* < 10^−4^), i.e., when *meg-3/4* hermaphrodites were provided with functional sperm. These results imply that the reduction in brood size in selfing *meg-3/4* hermaphrodites is at least in part due to defects in self-produced sperm. The three other small RNA mutants that we found to be prematurely attractive exhibit sperm defects such as abnormal sperm pseudopod formation(*50*, *56*) or shorter spermatogenesis(*51*). Collectively, these findings suggest that disruption of germ granules and mistargeting of sperm genes by small RNAs may cause mild sperm defects that trigger premature attraction. Indeed, as detailed above, severe defects in sperm are known to trigger the secretion of the male-attracting volatile pheromone(*7*, *8*).

**Fig.3.**
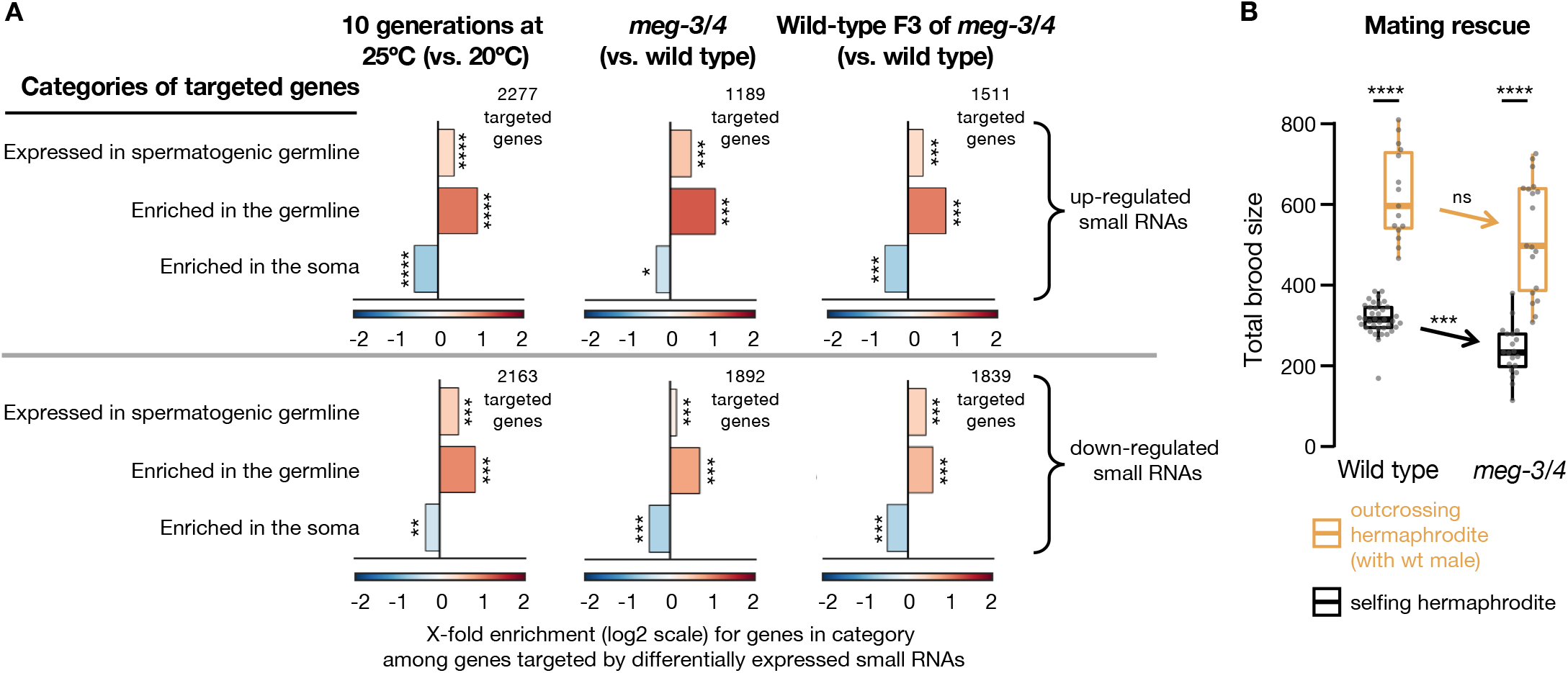
Prematurely attractive hermaphrodites exhibit misregulation of sperm genes and sperm defects. (**A**) Enrichment analysis of small RNAs up-regulated (high panel) and down-regulated (low panel) in worms upon cultivation at 25°C(*32*) (left), in *meg-3/4* mutants(*43*) (center) or in F3 wild-type progeny of *meg-3/4* mutants(*43*) (right). All small RNAs targeting a specific protein-coding gene were grouped together into one value that was used to assess whether the total amount of small RNAs targeting the gene was different (analysed with Deseq2, adjusted *P* value < 0.1). The number of differentially targeted protein-coding genes for each condition appears next to the relevant panel. Shown are fold (log2 scale) enrichment results (observed/expected) for genes known to be expressed in the spermatogenic germline(*67*), for genes enriched in the whole germline(*68*) and for genes enriched in the soma(*68*), in the lists of genes differentially targeted by small RNAs. Adjusted *P* values were obtained using a randomization bootstrapping test and corrected for multiple comparisons using the Benjamini–Hochberg step-up procedure. (**B**) Total brood size (y-axis) of wild-type and *meg-3/4* hermaphrodites reproducing via selfing (black) or via outcrossing (yellow). Insemination by wild-type males similarly increased the brood size of wild-type and *meg-3/4* hermaphrodites, indicating that *meg-3/4* oocytes can be fertilized when provided with wild-type sperm. Data collected over 3 independent experiments. Dots represent values for individual hermaphrodites. Boxplot (Tukey’s style): median & IQR, whiskers extend to the most extreme value within 1.5xIQR from the 25^th^ or 75^th^ percentile. Welch’s ANOVA, Games-Howell post-hoc correction for multiple comparisons. \*\*\*\**P* < 10^−4^, \*\*\**P* < 0.001, \*\**P* < 0.01, \**P* < 0.05, ns *P* > 0.05.

### Transgenerational premature attractiveness can affect the genetic structure of the population

Secreting a male-attracting pheromone could be costly (due to the costs of the production of the pheromone itself or the consequences of the mating it facilitates) and may thus negatively impact the fitness of hermaphrodites that have the “cheaper” option of reproducing via selfing(*57–59*). We used a theoretical population genetics approach to examine computationally, based on new empirical data that we collected (Fig. S10B and chemotaxis results), under which conditions a transient increase in premature male-attraction would spread in the population (see Methods for details). For secretion of the male-attracting pheromone to be adaptive our model predicted two requirements: (1) the stress should persist for multiple generations (2) there should be a mild increase in the number of males (Fig. 4A). The first condition is in line with our abovementioned observation that long-term, but not short-term, cultivation at 25°C induces premature attraction (Fig. 1A & Fig. S1). Interestingly, a recent study has found that *C. remanei* nematodes cope relatively better in elevated temperatures only if temperatures in previous generations were stably high or were gradually increasing in a “predictable” manner(*60*). In accordance with the second prediction of the model, <u>**h**</u>igh <u>**i**</u>ncidence of <u>**m**</u>ales (Him phenotype) was documented in worms grown at elevated temperatures (Fig. 4B and(*61*)). The Him phenotype also accompanies the transgenerational gradual loss of fertility (Mrt Phenotype) of *C. elegans* wild isolates and mutants impaired in small RNA and chromatin modification pathways when cultivated at high temperatures(*62*) (& reviewed in(*22*)). We discovered that selfing *meg-3/4* hermaphrodites and their wild-type descendants also have more male progeny than wild-type animals (Fig. 4C,D).

**Fig.4.**
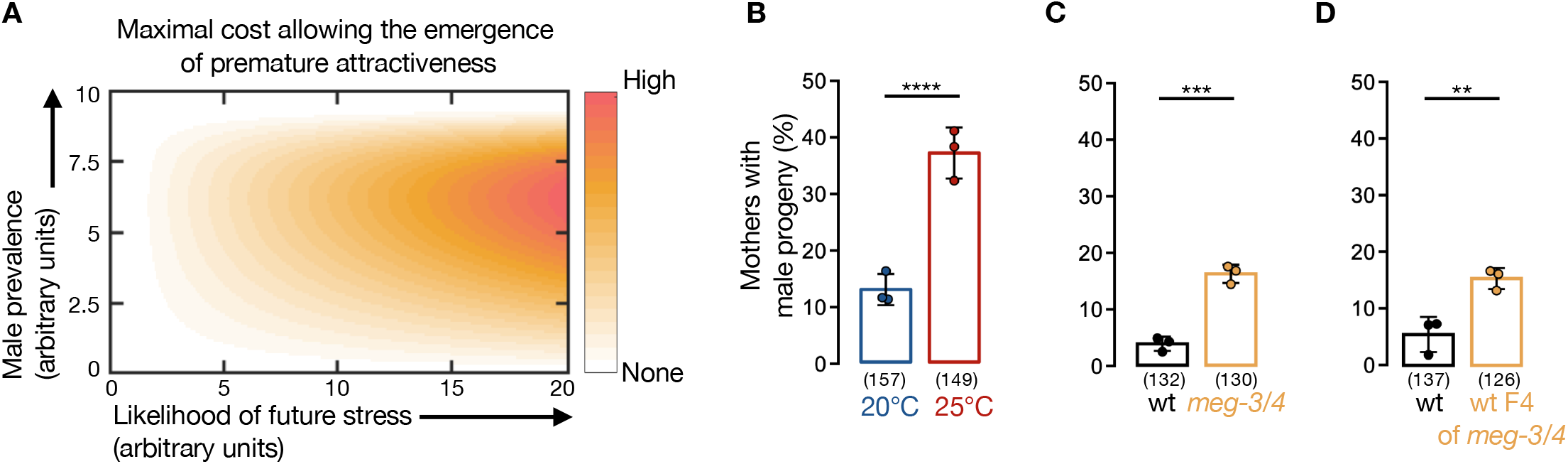
Premature attractiveness is associated with an increase in male prevalence. (**A**) Model for maximal cost (color-coded) that allows the evolution of premature attractiveness as a function of male prevalence in the population (y-axis) and likelihood of stress in future generations (x-axis). Model derivation and parameters are found in Methods and Supplemental Information. (**B-D**) Proportion (%) of hermaphrodites with males in their selfing-derived progeny. Tested conditions are indicated below data. Animals in (**B**) were wild-type N2 worms, while in (**C** & **D**) all animals bear an integrated single-copy *mex-5p::gfp* transgene in their genetic background. Dots represent results from independent experiments. Bars: mean ± sd. Numbers below bars: total *N* of tested hermaphrodites. Fisher’s exact test, \*\*\*\**P* < 10^−4^, \*\*\**P* < 0.001, \*\**P* < 0.01.

To examine whether heritable small RNA-mediated premature attractiveness can impact the genetic structure of a population, we conducted multigenerational competition experiments (scheme in Fig. 5A and Methods). In these experiments, we took advantage of the fact that premature attractiveness is strongly transmitted to wild-type descendants of *meg-3/4* mutants, and let worms which are genetically identical (wild type) but differ in their epigenetic heritage (naïve vs *meg-3/4* descendants) compete over the same resources (food and males). To determine whether the effects of the epigenetic heritage on the population are due to the male-attracting pheromone, we ran the experiments side-by-side using both wild types and *srd-1(−)* mutants that cannot sense the secreted pheromone.

**Fig.5.**
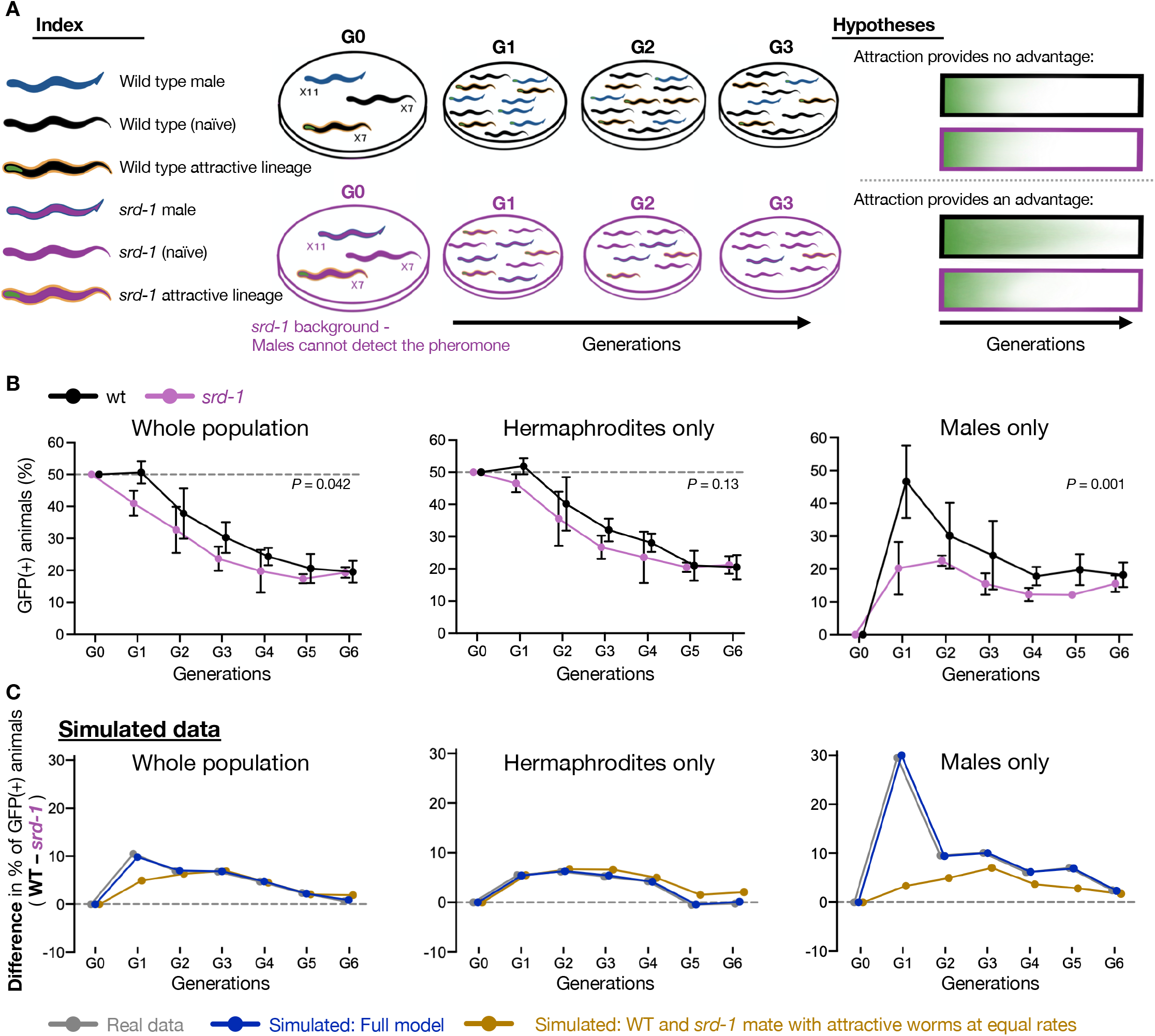
Transgenerational premature attractiveness can affect the genetic structure of the population. (**A**) Scheme depicting the multigenerational competition experiments. In these experiments, GFP-tagged wild-type animals originating from *meg-3/4(−)* ancestry (“attractive lineage”) competed against *naïve* GFP-negative wild-type animals (non-attractive lineage). Large populations were imaged and analysed in every generation. Experiments were performed side-by-side using two genetic backgrounds: wild-type (black) and *srd-1* mutants (purple). (**B**) Epigenetically attractive lineages mate more frequently and are more prevalent in populations when males sense the secreted male-attracting pheromone. Proportion (%) of GFP-positive animals (y-axis) over generations (x-axis) in whole populations (left) or sexual subgroups (center & right). Males present in the populations likely derive from outcrossing events, as outcrossing progeny contains 50% males vs only ~0.1% in selfing-derived progeny. Experiments with wild-type (black) and *srd-1(−)* (purple) animals were performed side-by-side, in three biological replicates. Dots and bars: mean ± sd. N per group/replicate/generation = 368±93, range 184-576. Indicated *P* values were calculated using the Generalized Linear Mixed Model (GLMM). (**C**) The difference (wild-type – *srd-1*) in the proportions (%) of GFP-positive animals (y-axis) over generations (x-axis). Shown are the experimental results (grey, as depicted in [**B**]), the simulated results (dark blue) and the simulated results where the underlying assumption that wild-type and *srd-1(−)* mate differentially with attractive hermaphrodites is relaxed (light blue). See Methods and Fig. S11 for full details about the models and additional tested assumptions.

As described earlier, the ‘attractive’ wild-type descendants of *meg-3/4* produce a ~15% lower brood size compared to naïve wild-type worms (Fig. S10A), and are thus expected to decrease in frequency across generations. Remarkably, we found that the ability of the males to sense the male-attracting odor increased the relative frequency of the ‘attractive lineage’ (compare *srd-1*(+) and *srd-1*(−) populations, Fig. 5B). Moreover, the prevalence of males deriving from the ‘attractive lineage’ was substantially higher in the population with functional SRD-1, indicating an increase in mating events. We also conducted *in silico* simulations of population dynamics incorporating empirical data that we collected on differences in male attraction, brood size, and lineage fitness (see Fig. S11 and Methods for more details). These analyses corroborated that the increase in male prevalence is largely explained by *srd-1*-dependent premature attractiveness (Fig. 5C & S11). Together, these results imply that detection of the male-attracting pheromone transgenerationally increases the mating frequency of prematurely attractive hermaphrodites, and the spread of alleles derived from heritably attractive lineages in the population.

## Discussion

Overall, our findings reveal that in response to mildly stressful temperatures, *C. elegans* hermaphrodites elevate their attractiveness and the attractiveness of their stress-naïve descendants for multiple generations. Mechanistically, premature attractiveness is associated with sperm defects and with disturbances in germ granules, and is transgenerationally inherited to ensuing generations via heritable small RNAs and the germline Argonaute HRDE-1. Since increased premature attractiveness also elevates the frequency of mating events, we conclude that transient small RNA-based responses to environmental challenges can impact genetic variation and could thus potentially shape the course of evolution. This phenomenon constitutes a novel manifestation of stress-induced evolvability, analogous to stress-induced mutagenesis, stress-triggered mobile elements transposition, horizontal gene transfer, and others(*63–66*). The heat-induced increase in evolvability that we describe here differs from these examples since it is driven by an inherited entity. Thus, the heritable small RNAs can affect genetic material by altering sexual behavior even when the original environmental stimulus is no longer present.

In addition to increasing *genetic* variability, we speculate that the inherited enhanced attractiveness and mating may facilitate the elimination of accumulated detrimental *epigenetic* information that could harm the animal’s reproductive fitness. Indeed, many mutants which are defective in small RNA inheritance exhibit a Mrt phenotype and become gradually sterile if forced to self-fertilize in the lab (reviewed in(*22*)). High temperatures impact sperm function in a wide range of species including mammals. In *C. elegans* hermaphrodites, elevated temperatures that induce sperm defects also trigger a heritable increase in sexual attraction that could be adaptive under environmental stress. Deepening our understanding of biological strategies to better cope with high temperatures is critical for the preservation of biodiversity amid global warming.

## Acknowledgements

We thank all members of the Rechavi lab and Shaham lab for helpful discussions. We would like to thank Daniel Leighton, Paul Sternberg, Gillian Stanfield and Ehud Cohen for their experimental advice, and Annie Mais for her help with data analysis. We thank Anat Nitzan, Ronen Zaidel-Bar and the Rockefeller University Bio-Imaging Resource Center for their expertise and assistance with imaging experiments. We thank Guy Teichman and Hila Gingold for their assistance with coding and with the bioinformatic analysis. Some strains were provided by the Caenorhabditis Genetics Center which is funded by the N.I.H. Office of Research Infrastructure Programs (P40 OD010440), and we thank Eric Miska for providing the SX1263 strain. SS is supported by N.I.H. grant R35NS105094. OR is thankful to the Adelis foundation grant #0604916191, the D.F.G. grant SCHU 2494/10-1, and E.R.C. grants #335624 & #819151 for funding.

## Author Contributions

IAT, IL, YM & OR conceived the project and designed the experiments. IAT, IL & YM performed the experiments and analysed the data with the assistance of YG & DF. YG & LH designed and implemented the mathematical model and the computer simulations. LHZ & OA generated the aid::ha::hrde-1 strains. SS provided guidance and resources. IAT, IL, YM & OR wrote the manuscript with input from all the authors.

## Methods

### Culture of worms and genetic crosses

All strains used in this study are listed in Key Resource Table. Worms were cultured using standard procedures in NGM plates seeded with *Escherichia coli* OP-50 and kept at 20°C unless noted otherwise. For the genetic crosses depicted in Fig. 2, S5,S7 & S10, 3-4 L4 hermaphrodites were placed with 10-15 males of the indicated genotypes in plates previously seeded with one drop of OP50. P0 Mated hermaphrodites were transferred to fresh plates without males after 16-24 hours of interaction. F1 hermaphrodites were separated to fresh plates as L4s to prevent mating by F1 males. Genotyping of F1s and F2s was performed by PCR.

### Extraction of Worm-Conditioned Media (WCM)

WCM (containing the odors secreted by the hermaphrodites) was produced as previously described(*7*) with minor modifications. To obtain large synchronized populations of hermaphrodites on the first day of adulthood, 12-20 gravid adults were transferred to fresh seeded plates and were allowed to lay eggs for 2-3 hours before removal. This procedure was performed in triplicates. 72-80 hours later (or 60-64h for animals grown at 25°C due to faster development) the adults were rinsed off the plates and washed 4 times with M9 buffer in 1.7ml tubes to remove bacteria. In every round of wash, worms were allowed to settle at the bottom of the tube for 90-120 seconds, then we removed the supernatant, applied 1ml of fresh M9 and gently mixed the tube via finger-tapping. After the last removal of the supernatant, worms were transferred to a non-seeded NGM plate with no cover using a Pasteur pipette. Once the buffer fully evaporated, 75 worms were transferred by picking into the inverted cap of a standard 1.7ml tube containing 75μl of M9. Caps were then sealed with parafilm and maintained for 24 hours at 20°C. Following incubation, the full contents of the caps were transferred into 1.7ml tubes and rested on ice for 3 minutes to let the worms and eggs settle at the bottom of the tube. 50μl of supernatant <u>with no worms/eggs</u> were then transferred into a new tube and stored at −20°C. WCMs from wild-type young hermaphrodites grown at 20°C were collected side-by-side with WCMs from biological groups of interest and used as negative control in subsequent male chemotaxis experiments.

25%-30% of *meg-3/4* adults are sterile and noticeably bear empty uteri(*53*), thus special care was taken to select only fertile worms containing visible embryos for WCM production.

In some cases, we used an alternative synchronization protocol in which L4 hermaphrodites were picked to new plates 24 hours before beginning of WCM incubation. These cases included the obligatory-outcrossing strain *fog-2*, strains with pronounced developmental delays (*alg-1*, *prg-1*), or animals deriving from genetic crosses and whose parents required PCR-genotyping. Control groups were collected similarly side-by-side and WCM extraction was performed normally.

### Male Chemotaxis experiments

Male chemotaxis assays were inspired from *Leighton et. al.*(*7*) with a few modifications. To obtain large populations of wild-type or *srd-1(−)* males, ten mating plates were prepared, that included twelve L4 hermaphrodites and twenty young males and a small lawn of food. 24h later, mated hermaphrodites were transferred to fresh seeded plates without males. 3 days later (one day before chemotaxis experiment), 1600-2000 young males were separated from hermaphrodites by picking and transferred to fresh seeded plates for 24 hours. This isolation was meant to reduce the impact of recent mating on the chemotaxis performance. Chemotaxis plates (90mm diameter, 2% agar, 5 mM KP0_4_, 1mM CaCl_2_ and 1mM MgS0_4_) were poured on the day of male picking, let dry with no cover for 30 minutes, then with cover overnight at room temperature. On the day of experiment, remaining excess of humidity on the plates’ cover was removed with a Kimwipe. 1μl of 1M sodium azide was dropped on each far side of the petri dish. Right after application of sodium azide on plates, WCMs were taken out of −20°C storage to thaw at room temperature, and males were rinsed from plates and washed four times in M9 (no spinning, just gravity). Once washes were completed, ~60-120 males were pipetted to the center of a chemotaxis plate. While the males were still swimming in the excess M9, 10μl of WCM were pipetted to the bottom side of the lid (facing the dish), above the sodium azide spot, and 10μl of M9 above the opposite spot (see scheme Fig. 1A). Excess of M9 in the plate was then gently dried out of the males’ area using a kimwipe, and the plate was covered with the media-applied lid. Plates were sealed with parafilm and incubated overnight in the dark at 20°C. At the conclusion of the assay, males on each side of plates were quantified, and all males within the 3cm central “buffer” zone were ignored. Chemotaxis index was calculated as (#males at WCM−#males at M9)/(#males at WCM+#males at M9). All chemotaxis experiments included a negative control (plates testing WCM extracted from young wild types) and a positive control (plates testing WCM from *fog-2* worms). Each experiment typically included 3-5 tested groups (controls included) and 3-4 biological replicates (chemotaxis plates) per group. Independent chemotaxis experiments were performed on different days and tested independently-produced WCMs. All biological groups of WCMs were tested across at least three independent experiments, with the exception of some of the screened mutants shown in Fig. 2A that did not trigger male-attraction.

### Exposure to environmental stress

#### L1 starvation

Synchronized gravid adults were bleached with standard hypochlorite treatment to obtain embryos and to remove traces of food. Embryos were transferred to NGM plates with no food, while a portion was transferred to OP50-seeded plates (fed controls). After 6 days of starvation, OP50 was added to the starved plates. WCM from starved and fed animals were collected side-by-side on the first day of adulthood (synchronized by picking of L4s 24h prior to WCM production). Due to differences in developmental pace, the WCM of fed worms were extracted from the F1 progeny of the “bleached” embryos.

#### Dauer induction

Dauer induction was conducted as previously described(*53*). Wild-type gravid adults were bleached with standard hypochlorite treatment. Embryos were resuspended in 5ml of S-Complete buffer containing 1mg/ml HB101 *E. coli*, at a concentration of 5 embryos/μl (i.e. 25,000 embryos in total). Worms were cultured in a 25mL Erlenmeyer flask at 20°C and 180 rpm for 40-45 days. Worms were eventually rescued upon transfer to NGM plates seeded with OP50. WCM from post-dauer animals was produced on the first day of adulthood. Control fed worms were cultivated in similar conditions after bleach except a higher HB101 concentration (38mg/ml) and lower embryo concentration (1 embryo/μl). The control worms were grown in liquid for 48 hours (they are L3 larvae by this time), and transferred to seeded NGM plates at the same time as post-dauer worms.

#### Cultivation at 25°C

For all data displayed in Fig. 1, wild-type worms had been cultivated at 25°C for 10-15 generations before initiation of the experiment. WCM production started 60h-64h after synchronized egg-laying and the 24h-incubation was performed at 20°C, similar to all other WCMs tested. WCM from wild-type 20°C controls were produced side-by-side, using worms that were synchronized by egg-laying 8h-12h prior to the 25°C synchronization (due to differences in developmental time). For the experiment depicted in Fig. S1B, G1 is the first generation grown at 25°C, and collection of WCM was performed similarly to the above. For transgenerational experiments (Fig. 1D-E & S4), the depicted P0 generation is the last generation cultivated at 25°C. P0 young adults were transferred to 20°C for 2-3 hours, to lay the synchronized F1 embryos. From this moment on, the F1s and the consecutive generations continuously grew at 20°C.

### Mating choice experiments

In those experiments, eleven males with fluorescent sperm interacted for exactly one hour on a small patch of food with seven “test” hermaphrodites and seven “competitor” hermaphrodites, and the mated hermaphrodites were subsequently scored based on fluorescence in their spermathecae.

One day before mating assays, young males (larvae with visible male morphology) were transferred into NGM plates containing 2μM MitoTracker™ Red CMXRos (Invitrogen, M7512) and previously seeded with OP50. On the same day, L4 hermaphrodites from the strains to be tested were picked into fresh plates, to isolate synchronized populations. For all experiments, the “competitor” hermaphrodites belonged to the pseudo-wt BFF53 strain that bears a *myo-2p::gfp* single-copy integrated transgene. After overnight incubation, males were rinsed off the MitoTracker™ plates and washed four times in M9, then recovered for 90 minutes on regular NGM plates with OP50. Seven *myo-2p::gfp* hermaphrodites (“competitor”) and seven hermaphrodites of the strain of interest (“test”) were then transferred to the center of a mating 60mm NGM plate (seeded with 50μl of OP50 at the center of the plate, 24h prior to experiment). Eleven Mitotracker™-stained males were added to the mating plate already containing the fourteen hermaphrodites. All 25 worms interacted for exactly one hour, at the end of which all males were removed. The 14 remaining hermaphrodites were mounted on a microscope slide with a 2% agarose pad, at least 1h after removal of males, to allow enough time for mating-derived sperm to reach the spermatheca(*69*). Hermaphrodites were imaged with x10 magnification using a BX63 Olympus microscope (Tel Aviv University) or a TiE Nikon microscope (Bio-Imaging Resource Center, Rockefeller University). Images were used to quantify the events of successful mating in each group of hermaphrodites, based on the presence of MitoTracker™-positive sperm in the spermathecae. “Competitor” and “test” worms were distinguished based on GFP fluorescence in the pharynx. Groups were typically tested in biological quadruplicates, and side-by-side with plates testing for wild-type and *fog-2* controls on the same day. Independent experiments were performed on separate dates, and all biological groups were tested across at least three independent experiments. In two instances (one plate in Fig. 1C and one plate in Fig S2, bottom right panel), one hermaphrodite from the competitor group was lost during slide preparation. In these two cases, the depicted proportion value was calculated out a total of 6 instead of 7.

### Generation of *aid::hrde-1* strains

We used a previously described CRISPR strategy with hybrid dsDNA donors(*70*) to tag the N’ terminus of the *hrde-1* gene with *aid::ha*. crRNA sequence: CAUAAUUUUGUCGAGCAAGU. The crRNA and *aid::ha* DNA sequence (PCR amplified to generate the repair templates) were synthesized by IDT. The ribonucleoprotein mix was injected as described(*70*), into animals of the SX1263 strain bearing a germline-expressed *mex-5p::gfp* transgene. The obtained engineered strain (BFF68) was then crossed with the CA1199 strain(*46*) bearing a *sun-1p::tir-1* transgene. The resulting strain (BFF69) was used for conditional HRDE-1 depletion experiments. Experiments to validate the auxin-dependent depletion of HRDE-1 are depicted in Fig. S3.

### Conditional *hrde-1* depletion experiments

Auxin treatment was performed as previously described(*46*) using the natural auxin indole-3-acetic acid (IAA) purchased from Alfa Aesar (#A10566). Auxin-containing plates were prepared by adding auxin in ethanol (1mM final concentration) to freshly-autoclaved NGM medium cooled to 50°C, just prior to the pouring of the plates. Concentrated (5x) OP50 was used on auxin-containing plates due to slower growth of bacteria on auxin.

#### Transgenerational RNAi experiments

These experiments were performed to test BFF69 animals for auxin-dependent disruption of heritable RNAi, the classic phenotype of *hrde-1(−)* worms(*20*). HT115 *E. coli* bacteria that transcribe anti*-gfp* dsRNA (or empty-vector controls) were grown overnight in LB supplemented with 25 μg/ml Carbenicillin. Bacterial cultures were then seeded onto NGM plates containing IPTG (1mM), Carbenicillin (25 μg/ml) and auxin (1mM). Bacteria were seeded on auxin+RNAi plates 3-4 days before addition of worms since bacteria grow slower on auxin-containing plates. L4-stage larvae of the BFF69 (or control strains, all bearing a *mex-5p::gfp* transgene) were transferred to auxin+RNAi plates overnight. The next morning, the adults (10-12 per plate) were allowed to lay eggs for 2-3 hours on fresh auxin+RNAi plates for synchronization of the P0 generation. Three days later, P0 adult worms were transferred to auxin-containing NGM plates with OP50 (no RNAi) and were allowed to lay eggs for 2-3 hours, for synchronization of the F1 generation. Synchronizations proceeded similarly every three days on auxin-containing plates seeded with OP50 until the F3 generation. On the day of synchronization, adults were collected for imaging on microscope slides with 2% agarose pads and a drop of M9 with 2mM levamisole. Images were acquired using a TiE Nikon microscope equipped with a Andor Neo sCMOS camera (Bio-Imaging Resource Center, Rockefeller University) with x10 magnification. Images were used to quantify the GFP silencing in the germline. Analysis of GFP fluorescence was done using ImageJ, by manually defining in each worm the area of the proximal oocyte and three background regions. CTCF values were calculated in this manner: Integrated density values of the oocyte – (area of measured oocyte * mean fluorescence of background regions). We normalized the CTCF value to the average CTCF value obtained from photographs of control animals of the same genotype, generation and age which were fed on empty-vector control plates.

#### Western blot

Young hermaphrodites were washed 4 times with M9 buffer, resuspended in ice-cold RIPA buffer and homogenized using a Dounce homogenizer. Following 3 minutes centrifugation at 850g, supernatants were collected in a new eppendorf tube and the protein concentration was quantified with BCA (ThermoFisher, 23227). Proteins were supplemented with sample buffer (ThermoFisher, [LC2676] + 2M DTT) and boiled for 5 minutes, then run on polyacrylamide gels in Tris-glycine running buffer (25mM Tris, 192 mM glycine, 0.1% SDS, pH 8.3). The proteins were transferred onto a nitrocellulose membrane, blotted membranes were blocked with 5% milk in TBST for 30 minutes and probed overnight (4°C) with primary antibody (anti-β-actin [Sigma-Aldrich, A5441], anti-HA [Biolegend, 901501]). After 3 washes with TBST, membranes were probed with Horseradish peroxidase-conjugated secondary antibody (1h, Jackson ImmunoResearch, 715-035-150), and visualized using ECL substrate (ThermoFisher, 34580) in an Amersham Imager 600 (GE).

#### Transgenerational premature attractiveness experiments

BFF69 worms were cultivated at 25°C for 10-15 generations, and the last generation of worms grown at 25°C were considered the P0 generation. Once they reached the first day of adulthood, P0 hermaphrodites were transferred to 20°C auxin-containing NGM plates (or control NGM plates) for 90 minutes, then transferred again to new auxin/control plates for 2-3 hours to lay the synchronized F1 embryos. The 90 minutes pre-incubation was performed to assure that all the F1 progeny derive from HRDE-1-depleted P0 mothers. For the rest of the experiment, all worms continuously grew at 20°C on auxin-containing/control plates, and WCM was extracted from young adults on every generation until the F3. WCM production and extraction was performed as described above and did not involve exposure to auxin. WCM from all conditions tested were collected side by side and appear in Fig. S4. The control BFF68 strain (that does not express the TIR-1 enzyme needed for auxin-dependent degradation of AID-tagged proteins) was exposed to auxin already from the P0 generation (25°C) and in following generations (at 20°C).

### Imaging of germ granules in the adult germline

BFF44 hermaphrodites were synchronized by picking young L4s 20h (20°C) or 16h (25°C for ten generations) prior to imaging. Young adults (with only one line of embryos in their uterus) were mounted on a microscope slide with a 3% agarose pad. Animals were imaged at their cultivation temperature thanks to a controlled closed chamber. Images were acquired at x100 magnification using a Nikon Ti-2 eclipse microscope equipped with a 100X CFI Plan-Apo 1.45 NA objective (Nikon, Tokyo, Japan), a CSU-W1 spinning-disk confocal head (Yokogawa Corporation, Tokyo, Japan), a DPSS-Laser (Gataca, France) and a Prime95B sCMOS camera (Photometrics, Tucson, AZ), controlled by the MetaMorph software (Molecular Devices, Sunnyvale, CA). We photographed the germline of young adult worms using ∼80 z stacks per worm with three channels: Bright field, 488nm and 561nm. Image analysis was performed with ImageJ software using the 3D object counter plugin(*72*).

### Analysis of RNA-Seq data

We used previously published datasets for *prg-1(−)* germline small RNAs and mRNAs(*72*), *meg-3/4(−)* mRNAs(*48*), *meg-3/4(−)* germline small RNAs(*43*), and germline small RNAs from F3 wild-type progeny of *meg-3/4* (second generation of homozygosity)(*43*). We performed re-analysis of recently published small RNA and mRNA expression data extracted from worms grown at 25°C(*32*), Accession number GSE138220. The quality of the fastq files were assessed with FastQC(*73*). Adapters were cut using Cutadapt(*74*) based on the adapter sequences taken from the original paper. Reads that were not cut or that were either less than 15 bp long (small RNA) or 17 bp (mRNA) long were removed. We also applied a quality threshold of 30, similar to the original publication. We used the following parameters:

For small RNAs:
“cutadapt -q 30 -m 15 --discard-untrimmed -a TGGAATTCTCGGGTGCCAAGG”.
For mRNAs:
“cutadapt -q 30 -m 17 --nextseq-trim=20 --max-n 2 -a AGATCGGAAGAGCACACGTCTGAACTCCAGTCA”

Next, small RNA reads were mapped to the *C. elegans* genome (WS235) using Shortstack(*75*) allowing no mismatches; mRNA reads were aligned using Bowtie2(*76*). The mapped reads were counted using HTseq_count(*76*) using a .gff feature file retrieved from wormbase.org (version WBcel235). The following parameters were used:

For small RNAs:
“HTSeq.scripts.count–stranded=reverse–mode=intersection-nonempty input.sam Genes.gff.”
For mRNAs:
“HTSeq.scripts.count --mode=intersection-nonempty input.sam Genes.gtf”

Differential expression was analysed using DESeq2(*77*). *P*-adjusted value <0.1 was regarded as statistically significant.

### Gene enrichment analysis

To test for enrichment for different gene list we used the “enrichment” function of the RNAlysis package (https://github.com/GuyTeichman/RNAlysis). The calculated fold enrichment values denote the ratio between the observed representation of a specific gene set to the expected one based on all genes (in this study we focused on all protein coding genes). Adjusted *P* values were obtained using a randomization test (10,000 random genesets), with the formula *P* = (successes + 1)/(repeats + 1). The *P* values were corrected for multiple comparisons using the Benjamini–Hochberg step-up procedure. For more information see RNAlysis documentation. We performed the analysis on 3 gene sets: (A) 752 germline-enriched genes(*68*), (B) 672 soma-enriched genes(*68*), (C) 6936 genes expressed in the spermatogenic germline(*67*). For this list we also applied a threshold of at least 10 RPKM.

### Hierarchical Clustering of siRNA expression

Hierarchical clustering was done using the clustermap function of the Seaborn(*78*) python package. Log2 fold change values in siRNA expression were taken from the DESeq2 output files. Default parameters were used: the distance metric was euclidic and the linkage method was average.

### Brood size quantification for self-reproducing and outcrossed hermaphrodites

Wild-type and *meg-3/4* L3 larvae were transferred individually to seeded plates (Day 0). After 24h (Day 1), each adult hermaphrodite was transferred to a new seeded plate (50μl drop of OP-50) containing 4 wild-type males, which were removed after 2 hours of interaction to avoid detrimental effects on hermaphrodite fitness(*79*, *80*). The hermaphrodites were transferred to new plates every 24h for 8 days. For each plate the progeny were quantified 3 days after transfer of the hermaphrodite. Hermaphrodites with numerous males in their progeny were considered mated, and non-mated hermaphrodites were used as (self-reproducing) controls. Results in Fig. 3B were collected in three independent experiments and included 2-16 hermaphrodites/group/replicate. For the data collected for the mathematical model (Fig. S10B), similar experiments were performed using wild-type hermaphrodites and BFF53 males bearing a *myo-2p::gfp* transgene. Quantified progeny were scored for their sex and for their genetic background (selfing/cross progeny) based on *gfp* expression using a fluorescent dissection microscope. Data from hermaphrodites that died before Day 6 of adulthood were excluded from the analysis. The experiment was conducted independently twice (R1 N_mated_ = 15, R1 N_unmated_ = 24, R2 N_mated_ = 11, R2 N_unmated_ = 23). For the experiment depicted in Fig S10A, experiments were run similarly to the above but did not involve any interaction with males. The experiment was conducted independently three times, and included 6-13 hermaphrodites/group/replicate.

### Mathematical model

We constructed a theoretical population genetics model to examine the range of conditions in which premature attraction would be favored by natural selection. Code available at: https://github.com/hadanylab/attraction.

#### Model Outline

For simplicity, we analysed a discrete-time model and assumed very large populations (so that demographic stochasticity and random genetic drift can be neglected) at carrying capacity, nonoverlapping generations, and single locus haploid inheritance. We performed an invasion analysis, exploring the success of a rare allele for premature attraction in a population of worms that do not display premature attraction. We derived the conditions for increase from rarity of said allele.

#### Model parameters

| Parameter | Description | Value |
| --- | --- | --- |
| $p$ | frequency of prematurely attractive worms in the population | $0 \leq p \leq 1$ |
| $m_1, m_2$ | mating rate for attractive and nonattractive worms, respectively | $0 \leq m_1, m_2 \leq 1$ |
| $X_c$ | brood size (of cross-derived offspring only) for mated hermaphrodites | 450 |
| $X_{cs}$ | brood size (of selfing-derived offspring only) for mated hermaphrodites | 100 |
| $X_s$ | brood size for unmated hermaphrodites | 330 |
| $f_m$ | frequency of spontaneous males | 0.001 |
|  | frequency of males in the progeny of mated hermaphrodites | 0.5 |
| $Y_{cms}$ | fitness of cross-derived male, under stress | 1 |
| $Y_{cfs}$ | fitness of hermaphrodite from crossing, under stress | $1 - s_m$ |
| $Y_{ms}$ | fitness of “spontaneous” male, under stress | $1 - s_c$ |
| $Y_{fs}$ | fitness of hermaphrodite from selfing, under stress | $(1 - s_m)(1 - s_c)$ |
| $Y_{cm}$ | fitness of male from crossing, under no stress | 1 |
| $Y_{cf}$ | fitness of hermaphrodite from crossing, under no stress | 1 |
| $Y_m$ | fitness of “spontaneous” male, under no stress | 1 |
| $Y_f$ | fitness of hermaphrodite from selfing, under no stress | 1 |
| $p_m$ | fraction of males in the population | $0 \leq p_m \leq 0.5$ |
| $s_m$ | male relative fitness advantage under stress | 0.1 |
| $s_c$ | crossed progeny relative fitness advantage under stress | 0.4 |
| $E_s$ | likelihood of stress | 0-1 |
| $C$ | cost of premature attraction | $0 \leq C \leq 1$ |
| $V$ | level of premature attraction | $0 \leq V \leq 1$ |

#### Mating function

We define *V*, level of attraction, measured in arbitrary units between 0 (= no attraction) and 1 (=max possible attraction), corresponding to chemotaxis values. We adapted a function from (*81*), to describe the probability of mating for an hermaphrodite as a function of the frequency of males in the population (*p_m_*) and the level of attraction the hermaphrodites exude (*V*).

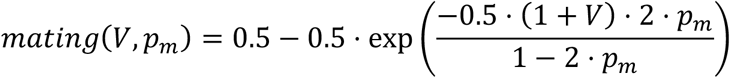

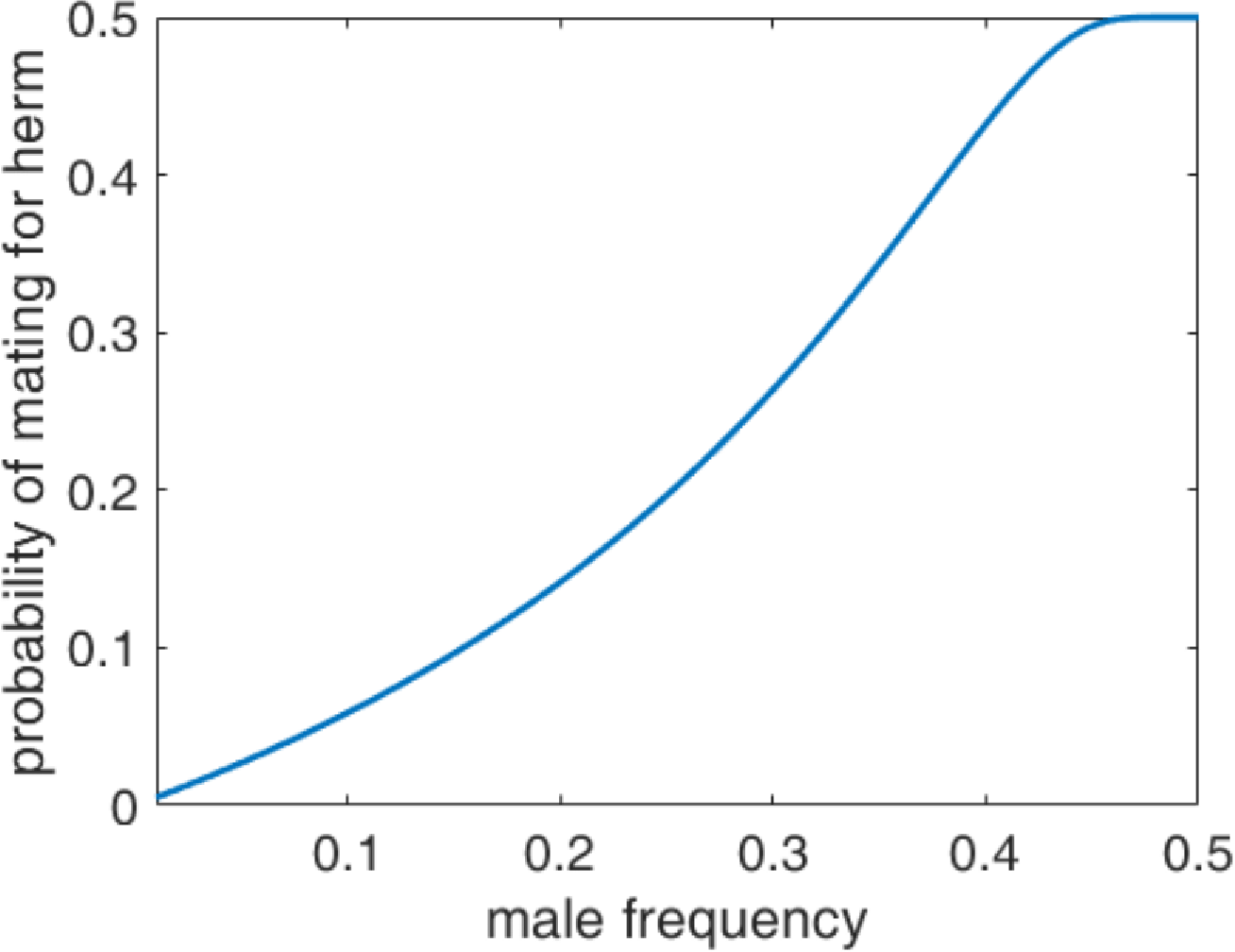
Shape of the mating function, parameters: *V* = 0

In the results presented in the manuscript, the values tested were *m*_1_ = *mating*(*V*_1_, *p_m_*) where *V*_1_ = 0.01, as we assumed in the invasion analysis that a rare variant will initially arise with a low level of attraction. For the rest of the population, *m*_2_ = *mating*(*V*_2_, *p_m_*) where *V*_2_ = 0, as we assumed the general population does not exhibit premature attraction.

#### Offspring types

We account for the possible offspring types that may exhibit the trait of premature attraction. We assume that males can carry and transmit the premature attraction trait to their offspring, but it does not affect their own phenotype.

In the following table, + marks an individual carrying the premature attraction trait, and – marks all other individuals.

| **Mother** | **Father** | **probability that offspring will be prematurely attractive** | **probability to form pairing of this type** | **brood size** |
| --- | --- | --- | --- | --- |
| + | + | 1 | $p \cdot (m_1 \cdot p)$ | $X_c$ |
| - | + | 0.5 | $p \cdot m_2 \cdot (1 - p)$ | $X_c$ |
| + | - | 0.5 | $(1 - p) \cdot (m_1 \cdot p)$ | $X_c$ |
| + | no father, unmated mothers | 1 | $p \cdot (1 - m_1)$ | $X_s$ |
| + | no father, selfing-derived progeny in mated mothers | 1 | $p \cdot m_1$ | $X_{cs}$ |

We adapted the different brood size numbers from the empirical data we gathered (Fig. S10B), as well as the relative fitness for every type of offspring. When there is no stress, we assume for simplicity that all types of offspring have the same fitness (thus, 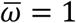). When there is stress, we assume that offspring born from outcrossing have an advantage (*s_c_*) over offspring born from selfing, and that males have an advantage (*s_m_*) over hermaphrodites.

#### Mean offspring fitness

Population mean mating probability:

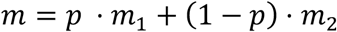

Total number of individuals in offspring generation (*N_H_* = number of hermaphrodites in the current generation):

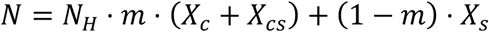

Mean offspring fitness without stress:

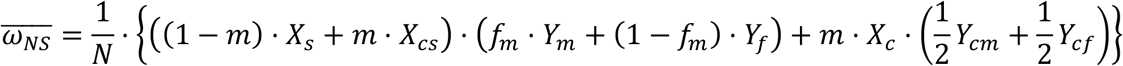

Mean offspring fitness under stress:

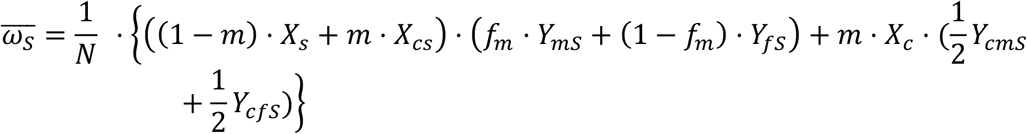

Mean fitness of individuals exhibiting premature attraction, in the following generation (without stress):

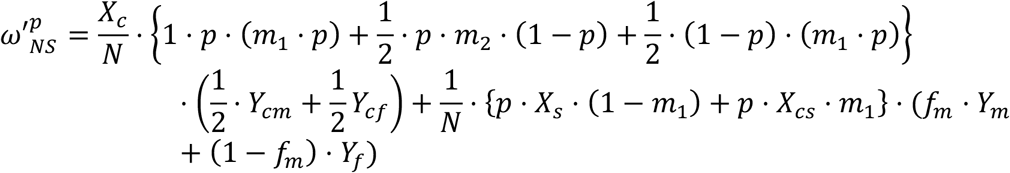

Mean fitness of individuals exhibiting premature attraction, in the following generation (under stress):

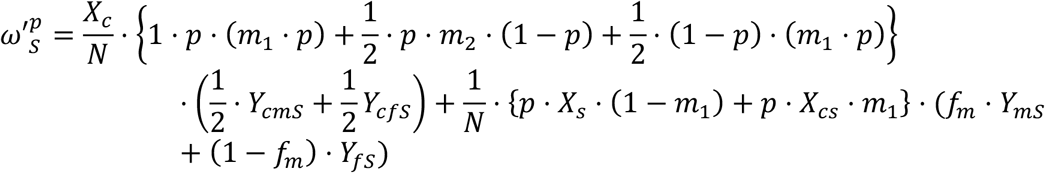

#### Evolution of premature attraction

We derive the frequency of individuals with premature attraction in the next generation, *p*′, given that in the current generation it is given by *p*. We separate the derivations into two scenarios, with stress and without stress.

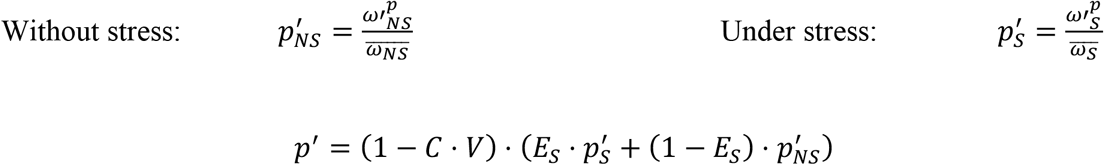

Where *C* is the cost of premature attraction, *V* is the level of premature attraction and *E_S_* is the likelihood of future stress.

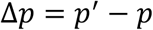

We are interested in the case of a rare variant of type 1 in a population of individuals of type 2. We want to find the condition for its increase from rarity, meaning 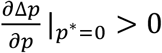. To do this, we find *C*\*, the critical *C* for which 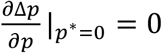. *C*\* is the maximal value for which 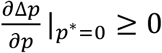., presented in Fig. 4A.

### Quantification of the incidence of males in selfing-derived progeny

Early L4 worms were transferred individually into 42-54 NGM wells (in standard 6-wells plates) seeded with 1-2 drops of OP50. Four days later (or 3.5 for worms grown at 25°C), when the progeny reached adulthood but food was still visible, the wells were examined for the presence of males using a dissection microscope. Each well was scanned for exactly 75 seconds and scored as positive if at least one male was detected during this time.

For experiments testing *meg-3/4* mutants (BFF40) and control wild types (SX1263), L4s were picked for synchronization, and individual young adults were individually transferred into the wells one day later to start the incubation period. This allowed us to include only fertile *meg-3/4* adults in the experiment.

### Multigeneration competition experiment

The “attractive” wild-type worms bore a *myo-2p::gfp* transgene and derived from outcrossed *myo-2p::gfp;meg-3/4* mutants, as described in scheme Fig. 2C. Seven “attractive” hermaphrodites (GFP+) and seven naïve wild-type hermaphrodites (GFP-negative) interacted 1h with wild-type males (GFP-negative) on a mating plate seeded with a dried 50μl drop of OP50. All worms were synchronized by picking L4s into a separate plate 16h prior to the 1h interaction. Side-by-side with the wild-type experiment, the same procedure was performed in a *srd-1(eh1)* background in which all worms involved (including for the ancestral crosses) carried the mutation. Both groups were tested in biological triplicates (a total of 6 groups). After 1h of interaction, all fourteen hermaphrodites (but not the males) were transferred together to fresh seeded plates (one per replicate), and were considered the P0 generation. **Step A**: After 40h, the worms were rinsed out of the plates into tubes, and allowed to settle for 2 minutes in a 1.7ml tube. Hundreds of F1 larvae were transferred into a new tube by pipetting the supernatant (by this time, the P0 adults reached the bottom of the tube). For each group, F1 larvae were transferred into seeded plates (~300-400 larvae per plate, in technical triplicates, 18 plates in total). **Step B**: After 56h, all F1s (now young adults) were collected, washed twice in M9, and transferred to fresh seeded plates in technical triplicates. This step permitted more interaction time for the adults without consuming all the food. At this occasion, 25 young males (GFP-negative, either wt or *srd-1* according to the group) were artificially added to each plate, representing 6%-8% of the cultured population. **Step C**: After 40h of incubation, all worms were rinsed out of plate using M9, F1 worms were kept for imaging (see below) and F2 larvae were separated like in Step A to form the next generation. Steps A-C repeated until the imaging of the F6 generation.

### Imaging and quantification of the multigenerational competition experiment

Collected adult worms (Step C of the above) were washed 8-10 times in M9 to eliminate the remaining food and larvae, with special care not to wash out the males that precipitate more slowly. After the last round of washes, worms were left in ~100μl and paralyzed in Sodium Azide (25-50mM final concentration). Worms were then transferred to imaging plates (60mm diameter, 8ml, 2% agarose, 0.3% NaCl, 5 mM KPO_4_, 1 mM CaCl_2_, 1 mM MgSO_4_) and physically separated from each other using a platinum-wired pick. Separation of the worms is important for allowing the *Wormachine* software to correctly process each worm individually(*82*). Images were acquired with x4 magnification using a TiE Nikon microscope equipped with a Andor Neo sCMOS camera (Bio-Imaging Resource Center, Rockefeller University). High throughput analysis of images was performed using the *Wormachine* software(*82*). Animals were binarily scored for GFP expression and for sex.

### Computer simulation of multigenerational competition experiment

We developed a computer simulation to reproduce the results of the multigenerational competition experiment under various settings (using Python version 3.7.9). The simulation receives as input the initial (G0) number of worms of each type: 7 *gfp(+/+)* hermaphrodites (attractive lineage), 7 *gfp(−/−)* hermaphrodites (naïve), 11 *gfp(−/−*) males. In every generation, the offspring distribution is calculated according to the following parameters: mating rates for hermaphrodites of the attractive lineage [*gfp(+/+)* or *gfp(+/−)*] or naïve lineage [*gfp(−/−)*], relative fitness (brood size) of attractive worms [i.e. *gfp(+/+)* or *gfp(+/−)*], brood sizes and the distribution of selfing-derived and cross-derived offspring. The amount of offspring for each genetic background combination is calculated. Then, 300 worms are sampled at random to form the next generation, and so forth.

Simulation parameters (derived from empirical results):

| Parameter | Value |
| --- | --- |
| brood size (of cross-derived offspring only) for mated hermaphrodites | 450 |
| brood size (of selfing-derived offspring only) for mated hermaphrodites | 100 |
| brood size for unmated hermaphrodites | 330 |
| frequency of spontaneous males | 0.001 |

We used manual parameter optimization to fit the unknown variables: mating rates and fitness of the green worms, such that the resulting dynamics aligns with the experimental data most accurately. Additionally, we tested different scenarios relaxing various underlying assumptions of the experiment. Full code available at: https://github.com/hadanylab/attraction.

Tested scenarios (see Fig. S11):

1. Full model predicting the real data
2. Fitness of the attractive lineage (*gfp*+) remains constant throughout the generations.
3. Attractive worms have the same mating rates in srd-1 and wild-type experiments.
4. Attractive and naïve worms are equal in fitness.
5. Attractive and naïve worms have the same mating rates.

### Statistical analyses

Statistical analyses and generation of graphs were performed using Prism, R v4.0.0 or MATLAB R2019b. Statistical details of experiments appear in figure legends. For the <u>chemotaxis experiments</u>, we applied the d’Agostino-Pearson omnibus test (α=0.05) to assess the normality of the values’ distribution in each tested group. In cases where assumption of normality was rejected or the test could not apply (N<8), we used the a-parametric Kruskal-Wallis test with Dunn’s correction for multiple comparisons to control. Otherwise, we used one-way ANOVA with Dunnett’s correction for multiple comparisons to control. For transgenerational experiments involving only two conditions, we used two-way ANOVA with Sidak’s correction for multiple comparisons. For the transgenerational HRDE-1-depletion experiment (involving five conditions), we used two-way ANOVA with Tukey post-hoc correction for multiple comparisons. For <u>the mating choice assays</u>, we used the Wilcoxon matched-pairs signed rank test (two-tailed). For <u>brood size and mating rescue experiments</u>, and because the assumption of equal variance was rejected (Brown-Forsythe test with *P* < 0.05) used Welch’s ANOVA with Games-Howell post-hoc correction for multiple comparisons. For <u>spontaneous male incidence</u> experiments, we used Fisher’s exact test. For the <u>multigenerational competition test</u>, we fit a generalized linear mixed model (GLMM) with generation, genetic background (wt vs. *srd-1*) and their interaction as fixed effects, using the using lme4 v1.1-21 package in R. Replicate populations were included as a random effect.

## Notes

### Competing Interest Statement

The authors have declared no competing interest.

