## Supplemental Information for "Transgenerational Regulation of Sexual Attractiveness in *C. elegans* Nematodes"

### Toker et al. – Supplemental Information

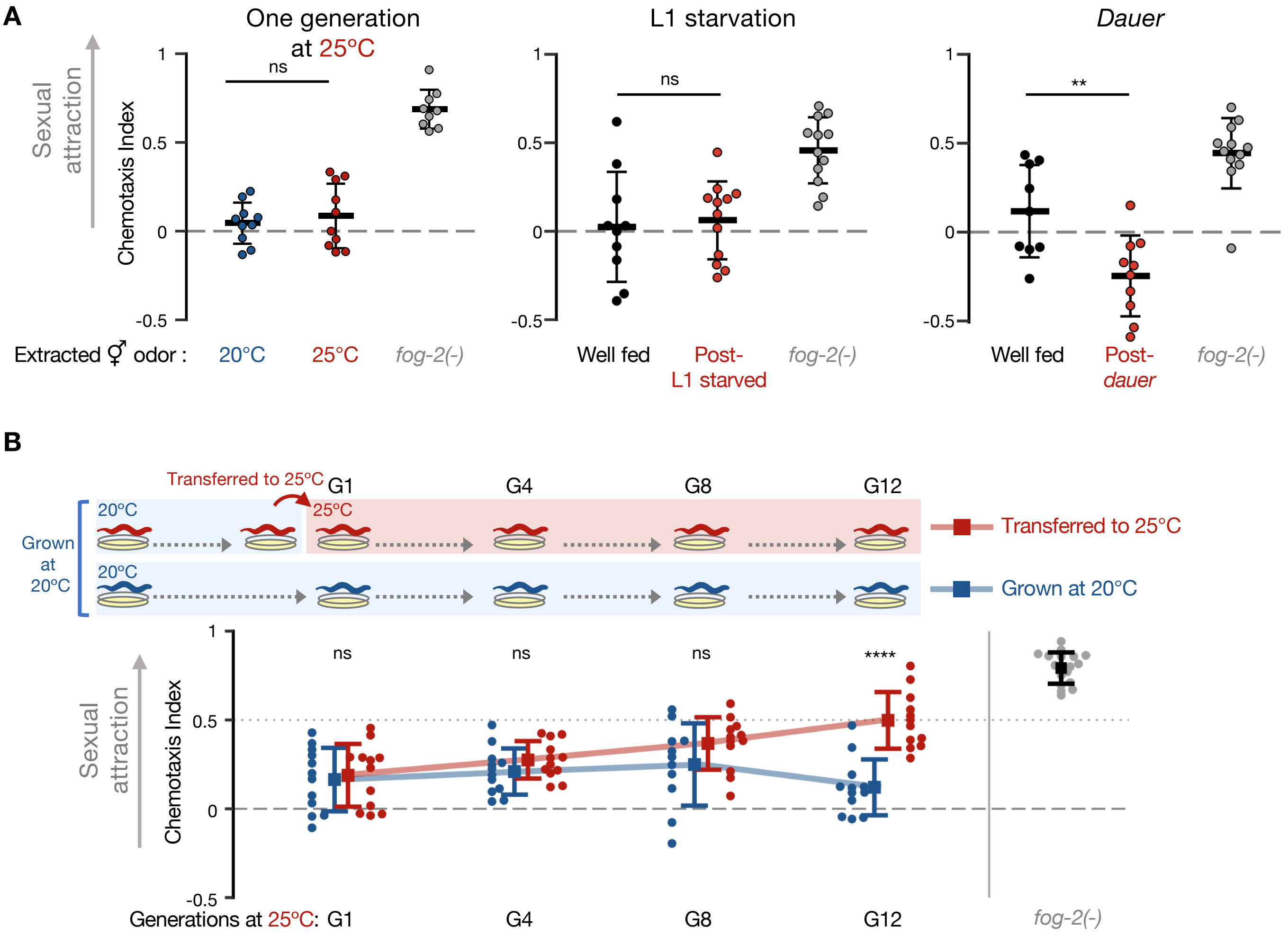


**Fig. S1. Examination of premature attraction among hermaphrodites exposed to different types of environmental stress. (A)** Male chemotaxis experiments. Tested odorants were extracted from one-day-adult hermaphrodites that were grown at 25ºC (left), that had been starved for 6 days at the L1 larval stage (center) or for 40-45 days in the alternative *dauer* stage (right). One-way ANOVA, Dunnett's correction for multiple comparisons to control. **(B)** Male chemotaxis experiments with odors extracted from hermaphrodites originally grown at 20ºC, transferred to 25ºC and tested after 1, 4, 8 and 12 generations at 25ºC (red). Continuously 20ºC-grown hermaphrodites were used as control (blue). Two-way ANOVA, Sidak's correction for multiple comparisons for every generation. **(A-B)** Each dot represents one biological replicate (chemotaxis plate) with 41-145 wild-type males. Bars: Mean ± sd, results from 3 independent experiments. ****$P < 10^{-4}$, **$P < 0.01$, ns $P > 0.05$.

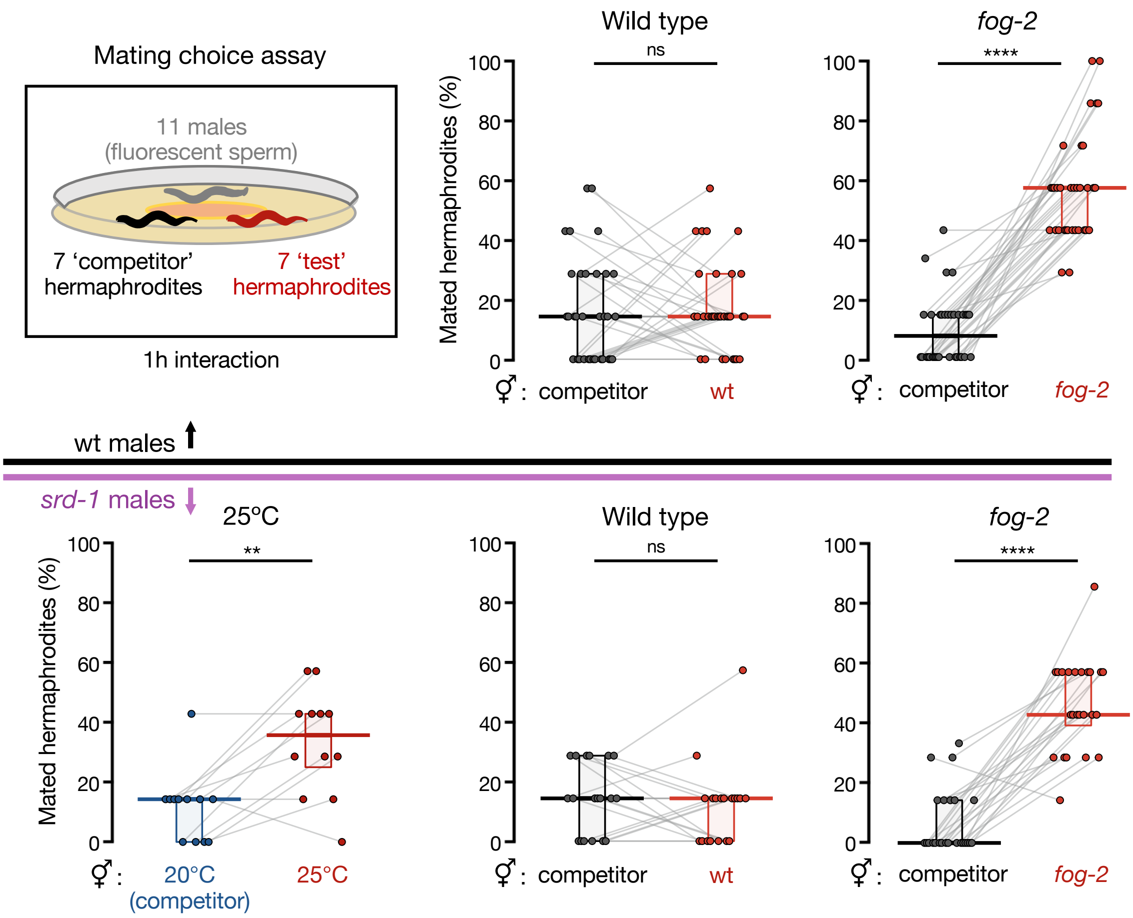


**Fig. S2**. **Additional mating choice experiments**. Eleven Mitotracker™-stained males interacted for exactly one hour with 14 young adult hermaphrodites divided evenly into two groups, "competitor" and "test". In all experiments, the "competitor" hermaphrodites contained an integrated single-copy transgene driving the expression of GFP in the pharynx (strain BFF53). Shown are the proportions (%) of mated hermaphrodites during the 1-hour interaction window, based on the presence of fluorescent sperm in their spermathecae. Experiments were performed using wild-type males (higher panel, controls for Fig. 1C) and *srd-1(-)* males (lower panel). Each grey line represents one biological replicate (mating plate), data collected from at least three independent experiments. Horizontal bar: Median. Boxes: Interquartile range. Two-tailed Wilcoxon matched-pairs signed rank test. $****P < 10^{-4}$, $**P < 0.01$, ns $P > 0.05$.

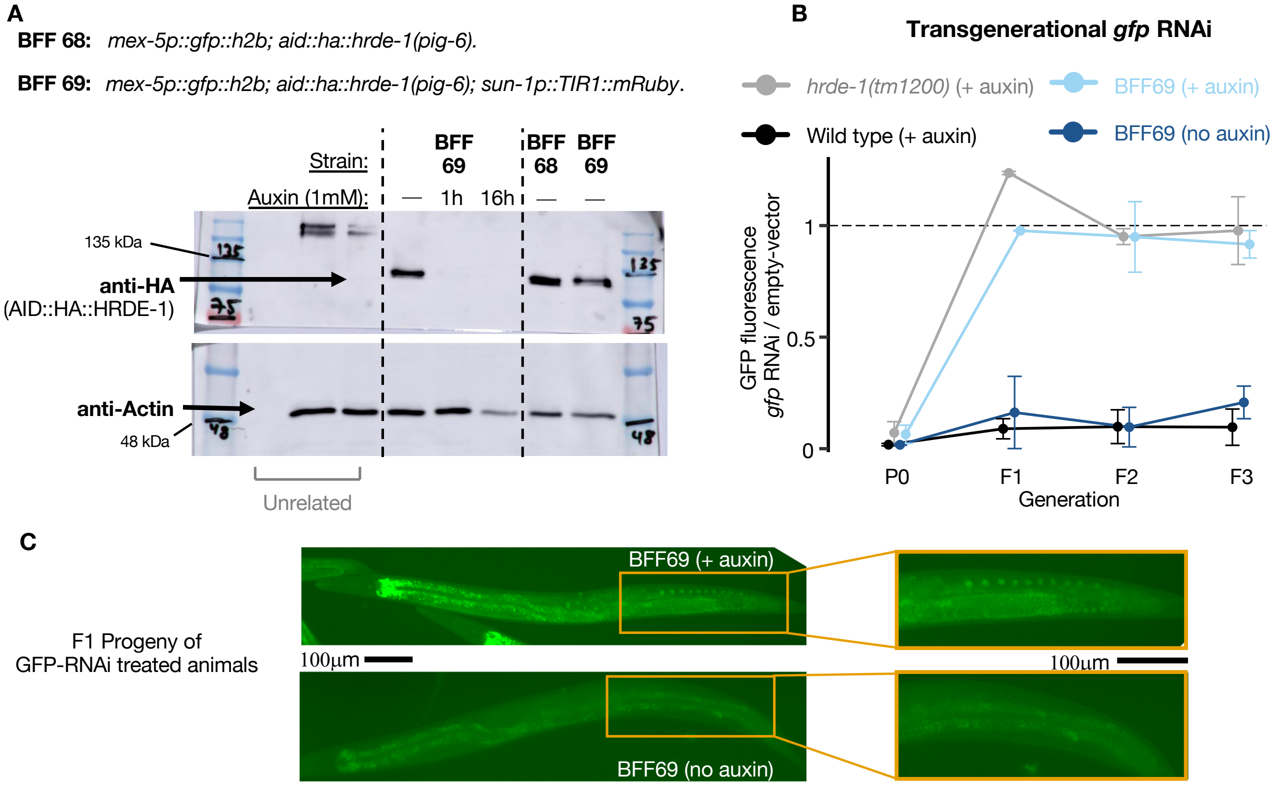


**Fig. S3**. **Conditional knockdown of HRDE-1 using the auxin-inducible degradation (AID) system.** In presence of the plant hormone auxin, the enzyme TIR-1 recognizes proteins containing the 44-amino acid AID tag and send them to degradation(*1*, *2*). Using CRISPR-Cas9, we inserted an *aid::ha* tag at the N' terminus of the endogenous *hrde-1* open reading frame, in worms carrying a single-copy transgene expressing GFP in the germline (BFF68), and subsequently crossed them with worms expressing the TIR-1 enzyme in the germline (BFF69). (**A**) Western blot analysis of lysates extracted from *aid::ha::hrde-1; sun-1p::tir-1* animals exposed to 1mM auxin. (**B**) Experiments testing for deficiency in heritable RNAi, the classic phenotype of *hrde-1* knock-out mutants(*3*). All tested strains contained a single-copy transgene driving the expression of GFP in the germline. To induce a transgenerational RNAi response in the germline, animals were cultivated on plates with bacteria expressing *gfp*-dsRNA (or empty-vector control) for one generation (P0), and transferred to plates with regular bacteria in following generations (F1-F3). GFP fluorescence (y-axis) is depicted relative to the mean fluorescence of control worms of the same genotype exposed to empty-vector control bacteria (at the P0 generation) and imaged side-by-side. Shown are mean ± sd from two independent experiments. N per group/replicate/generation = 58±14, range 31-88. (**C**) Representative images of *gfp* fluorescence in the germline of F1 progeny of *gfp*-RNAi-treated BFF69 worms (from experiment depicted in [**B**]).

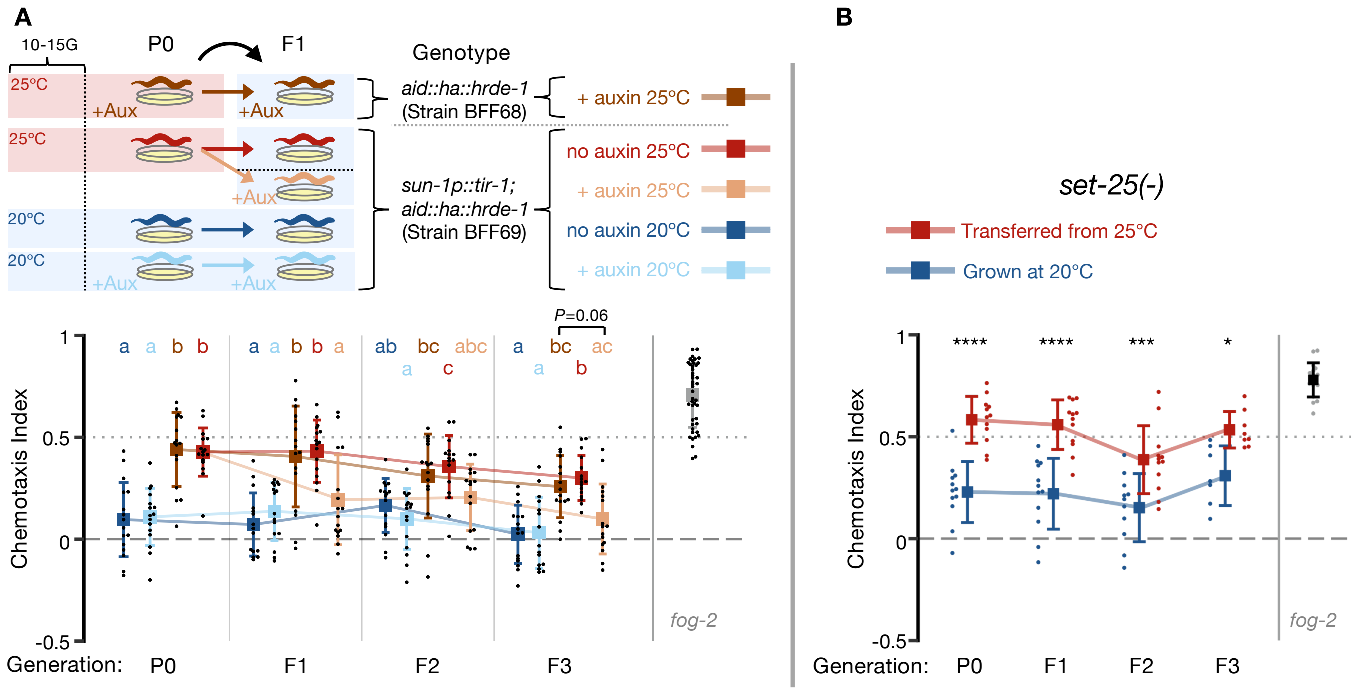


**Fig. S4**. **Transgenerational inheritance of premature sexual attraction requires the heritable small RNA machinery.** (**A**) Male chemotaxis experiments with odors extracted from *aid::hrde-1;sun-1p::tir-1* hermaphrodites grown at 25ºC for 10-15 generations (P0) and transferred back to 20ºC for 3 generations in the presence (beige, HRDE-1-depleted) or absence (red) of auxin. The data from Fig. 1E appears again here for convenience of visualization purposes, together with additional control groups. Extraction of odors, and male chemotaxis experiments testing them, were performed side by side for all depicted groups. Control groups included animals continuously grown at 20ºC with (light blue) or without auxin (dark blue), and *aid::hrde-1* animals lacking the *tir-1* gene (BFF68 strain) transferred from 25ºC in the presence of auxin (brown). Each dot represents one biological replicate (chemotaxis plate) with 38-138 males. Bars: mean ± sd, results from 4 independent experiments. Within each generation, data labelled with different letters are significantly different from each other ($P$<0.05), two-way ANOVA with Tukey's correction for multiple comparisons. (**B**) Male chemotaxis experiments with odors extracted from *set-25(-)* worms grown at 25ºC for 10-15 generations (P0) and transferred back to 20ºC for 3 generations (red), and *set-25(-)* worms continuously grown at 20ºC (blue). Each dot represents one biological replicate (chemotaxis plate) with 43-131 males. Bars: mean ± sd, results from 3 independent experiments, except for the F3 generation (two independent experiments). Two-way ANOVA, Sidak's correction for multiple comparisons for every generation. ****$P < 10^{-4}$, ***$P < 0.001$, *$P < 0.05$.
See also Figure 1.

| Genotype | Allele | Chemotaxis index (mean) | Replicates (N) | P value (vs. wt) |
|---|---|---|---|---|
| Wild type (N2) | – | 0.007 | 32 | – |
| *ppw-1* | *pk1425* | -0.225 | 3 | 0.9999 |
| *alg-1* | *gk214* | -0.087 | 4 | 0.9999 |
| *rsd-2* | *pk3307* | -0.049 | 3 | 0.9999 |
| *alg-2* | *ok304* | -0.003 | 4 | 0.9999 |
| *rde-4* | *ne299* | 0.050 | 18 | 0.9999 |
| *eri-1* | *mg366* | 0.051 | 5 | 0.9999 |
| *hrde-1* | *tm1200* | 0.070 | 3 | 0.9999 |
| *rde-11* | *hj37* | 0.147 | 8 | 0.9999 |
| *sid-1* | *qt9* | 0.206 | 19 | 0.5639 |
| *alg-3/4* | *ok1041 ; tm1155* | 0.208 | 3 | 0.9999 |
| *rrf-3* | *pk1426* | 0.261 | 3 | 0.9999 |
| *alg-5* | *tm1163* | 0.38 | 10 | 0.0074 |
| *dcr-1(Δhelicase)* | *mg375* | 0.556 | 4 | 0.0118 |
| *meg-3/4* | *tm4259 ; ax2026* | 0.569 | 3 | 0.0363 |
| *prg-1* | *n4357* | 0.772 | 3 | 0.0022 |
| *fog-2* | *q71* | 0.559 | 29 | $<10^{-4}$ |

**Table S1. Screen of small RNA processing mutants for premature attractiveness.** Information about the results of the screen depicted in Fig. 2A. All odors were extracted from hermaphrodites on the first day of adulthood. The *alg*-5 data appears also in Fig. S5. *P* values were obtained via Kruskal-Wallis test with Dunn's correction for multiple comparisons to wild type.

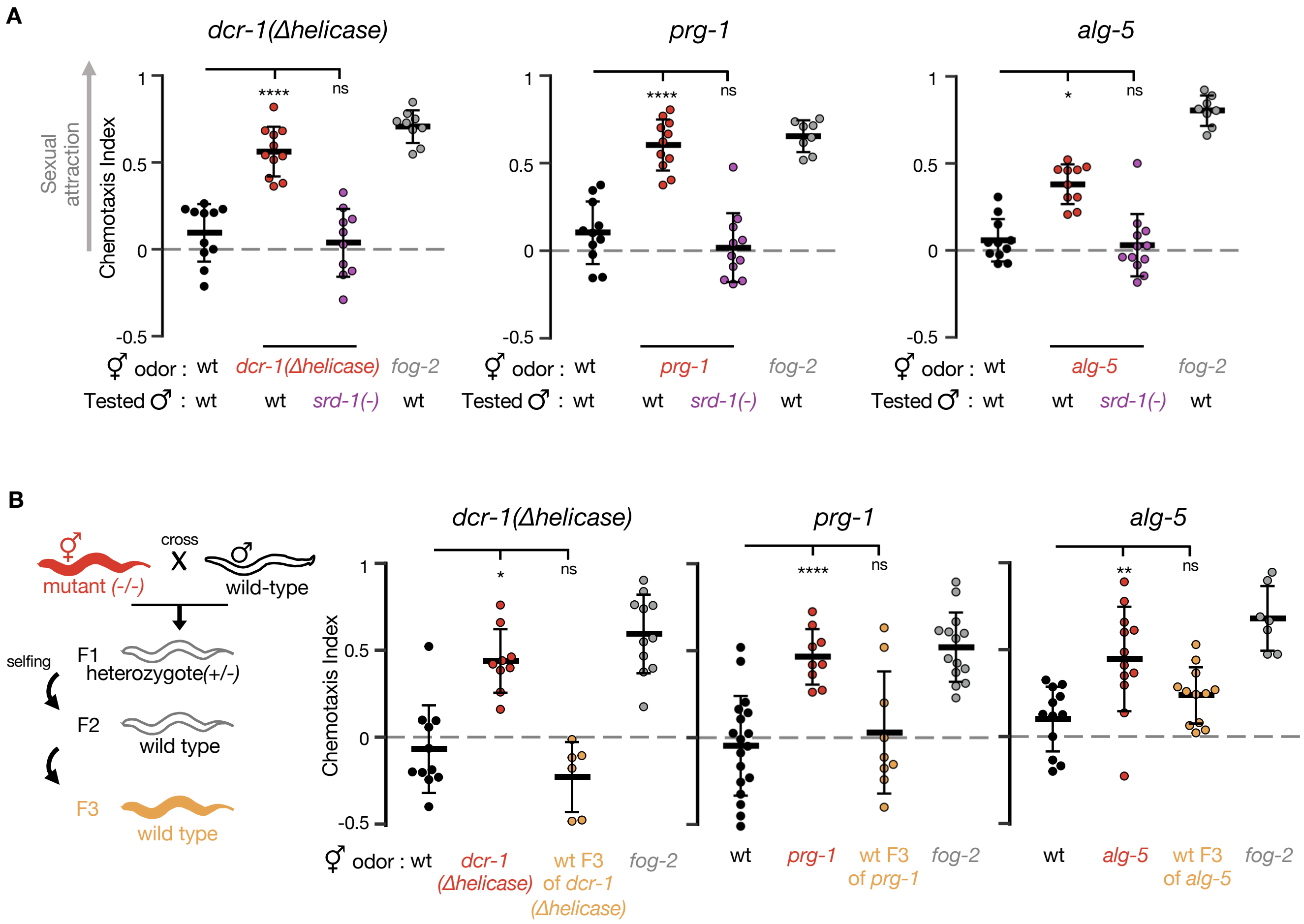


**Fig. S5**. **Hermaphrodites mutated in small RNA processing genes display premature attractiveness**. (**A**) Male chemotaxis experiments. Genotypes of hermaphrodites used for odor extraction and of tested males appear below panel. Each dot represents one biological replicate (chemotaxis plate) with 38~163 males. Some of the *alg-5* data appears also in Fig. 2A. One-way ANOVA, Dunnett's correction for multiple comparisons to wild-type (except for *alg-5* panel: Kruskal-Wallis test with Dunn's correction for multiple comparisons to wild-type).
(**B**) Male chemotaxis experiments testing for odors extracted from wild-type descendants of *dcr-1(Δhelicase), prg-1* & *alg-5* mutants. Each dot represents one biological replicate (chemotaxis plate) with 38-133 males. One-way ANOVA, Dunnett's correction for multiple comparisons to wild-type (except for *dcr-1(Δhelicase)* panel in [**B**]: Kruskal-Wallis test with Dunn's correction for multiple comparisons to wild-type). (**A-B**) Bars: mean ± sd. ****$P < 10^{-4}$, **$P < 0.01$, *$P < 0.05$, ns $P > 0.05$.

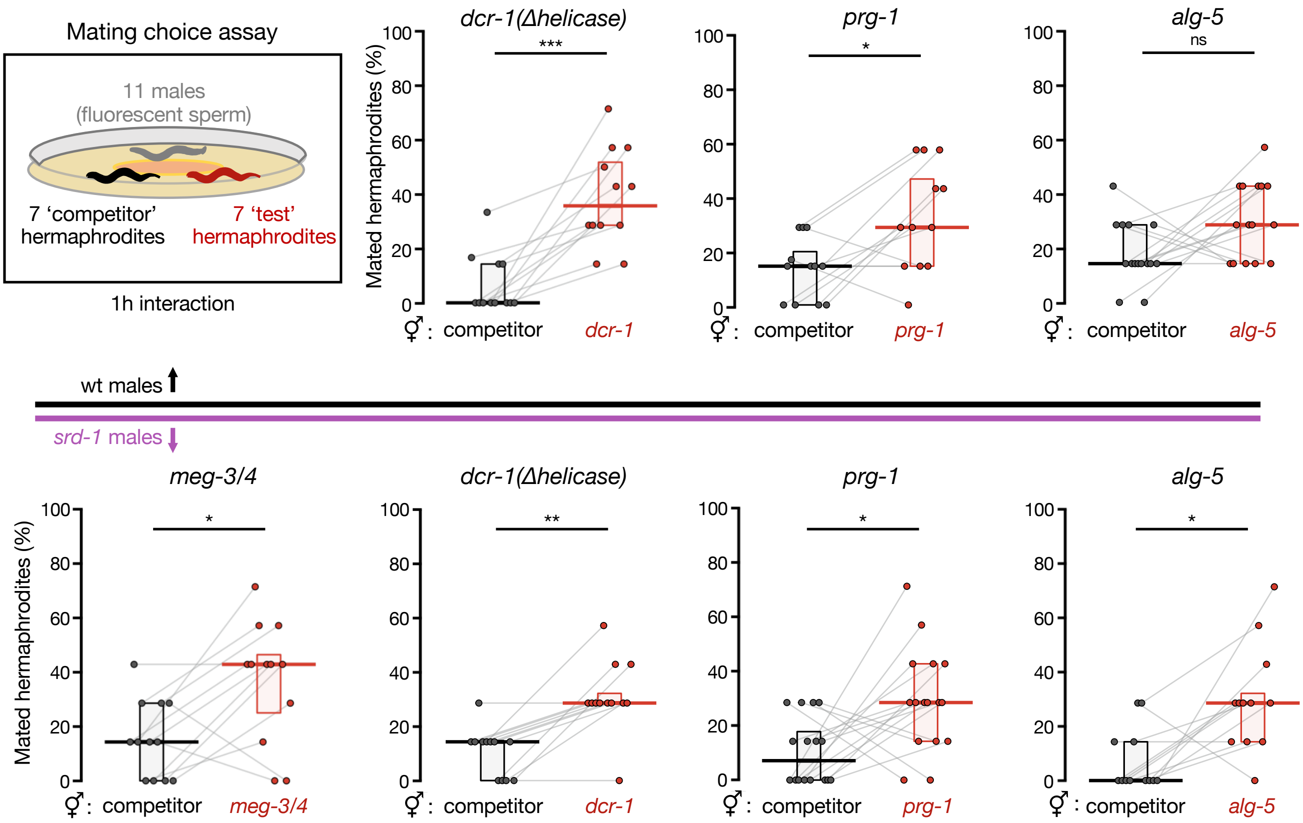


**Fig. S6. Prematurely attractive mutants outcross more.** Mating choice assays with prematurely attractive small RNA mutants. In all experiments, the "competitor" hermaphrodites had an integrated single-copy transgene driving the expression of GFP in the pharynx (strain BFF53). Shown are the proportions (%) of mated hermaphrodites during the 1-hour interaction window, based on the presence of fluorescent sperm in their spermathecae. Experiments were performed using wild-type males (higher panel) and *srd-1(-)* males (lower panel). Each grey line represents one biological replicate (mating plate), data collected from at least three independent experiments. Horizontal bar: Median. Boxes: Interquartile range. Two-tailed Wilcoxon matched-pairs signed rank test. ***$P < 0.001$, **$P < 0.01$, *$P < 0.05$, ns $P > 0.05$.

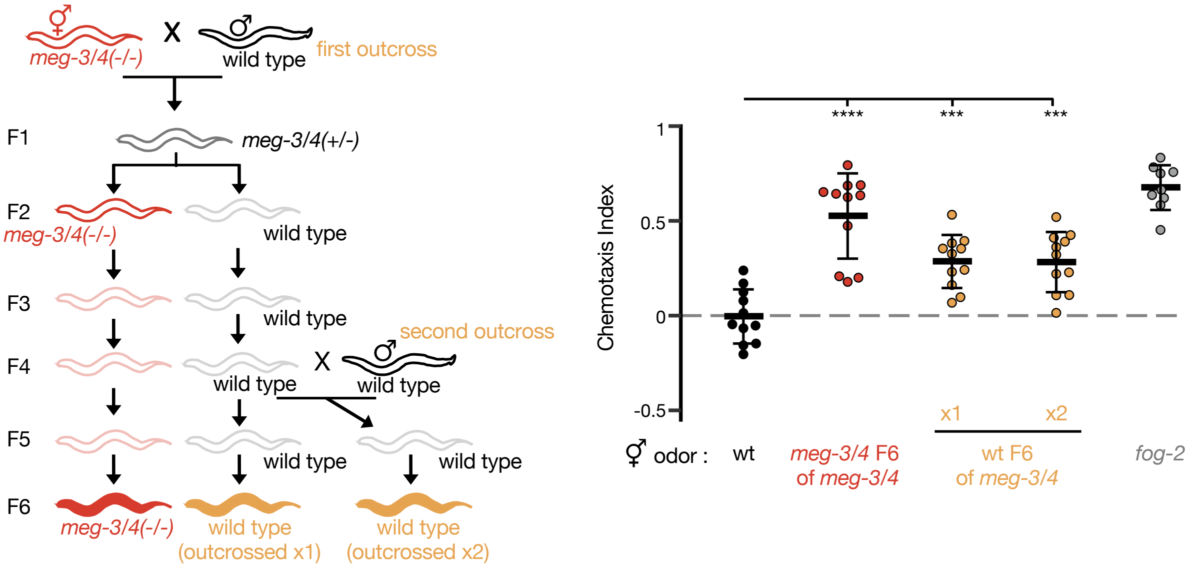


**Fig. S7. *meg-3/4* parents transgenerationally transmit premature attractiveness for six generations.** Male chemotaxis experiments testing for odors extracted from F6 wild-type descendants of *meg-3/4* double mutants outcrossed either once (P0) or twice (P0 & F4) with wild-type males. Each dot represents one biological replicate (chemotaxis plate) with 53~124 males. One-way ANOVA, Dunnett's correction for multiple comparisons to wild-type. Bars: mean ± sd. Data collected across 3 independent experiments. $^{****}P < 10^{-4}$, $^{***}P < 0.001$.

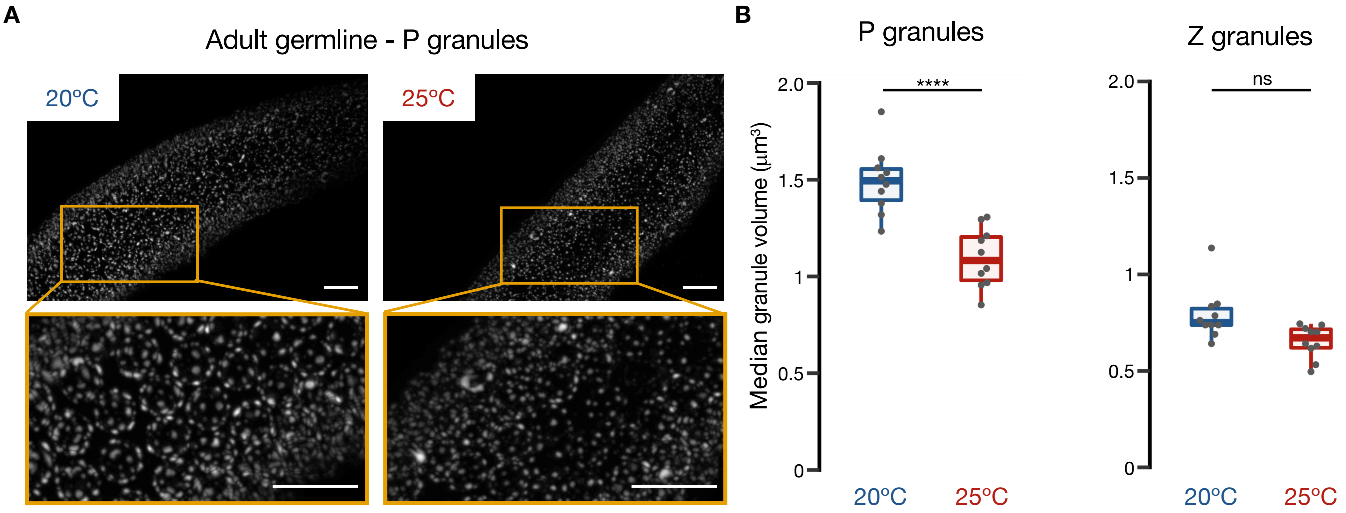


**Fig. S8. Hermaphrodites grown at 25ºC display smaller P granules.** (**A-B**) BFF44 hermaphrodites (N=10 per condition) bearing a *pgl-1::tagRFP* transgene (staining P granules) and a *wago-4::gfp* transgene (Z granules) were imaged as early young adults. (**A**) Representative micrographs of PGL-1::tagRFP fluorescence in the early pachytene zone of hermaphrodites grown at 20ºC (left) and 25ºC for ten generations (right). Scale bars: 10µm. (**B**) Median volume of germ granules in adults grown at 20ºC (blue) and 25ºC for ten generations (red). Each dot represents the median for one individual animal with >1000 analysed granules. Boxplot (Tukey's style): median & IQR, whiskers extend to the most extreme value within 1.5xIQR from the 25th or 75th percentile. One-way ANOVA, Sidak's correction for multiple comparisons. ****$P < 10^{-4}$, ns $P > 0.05$.

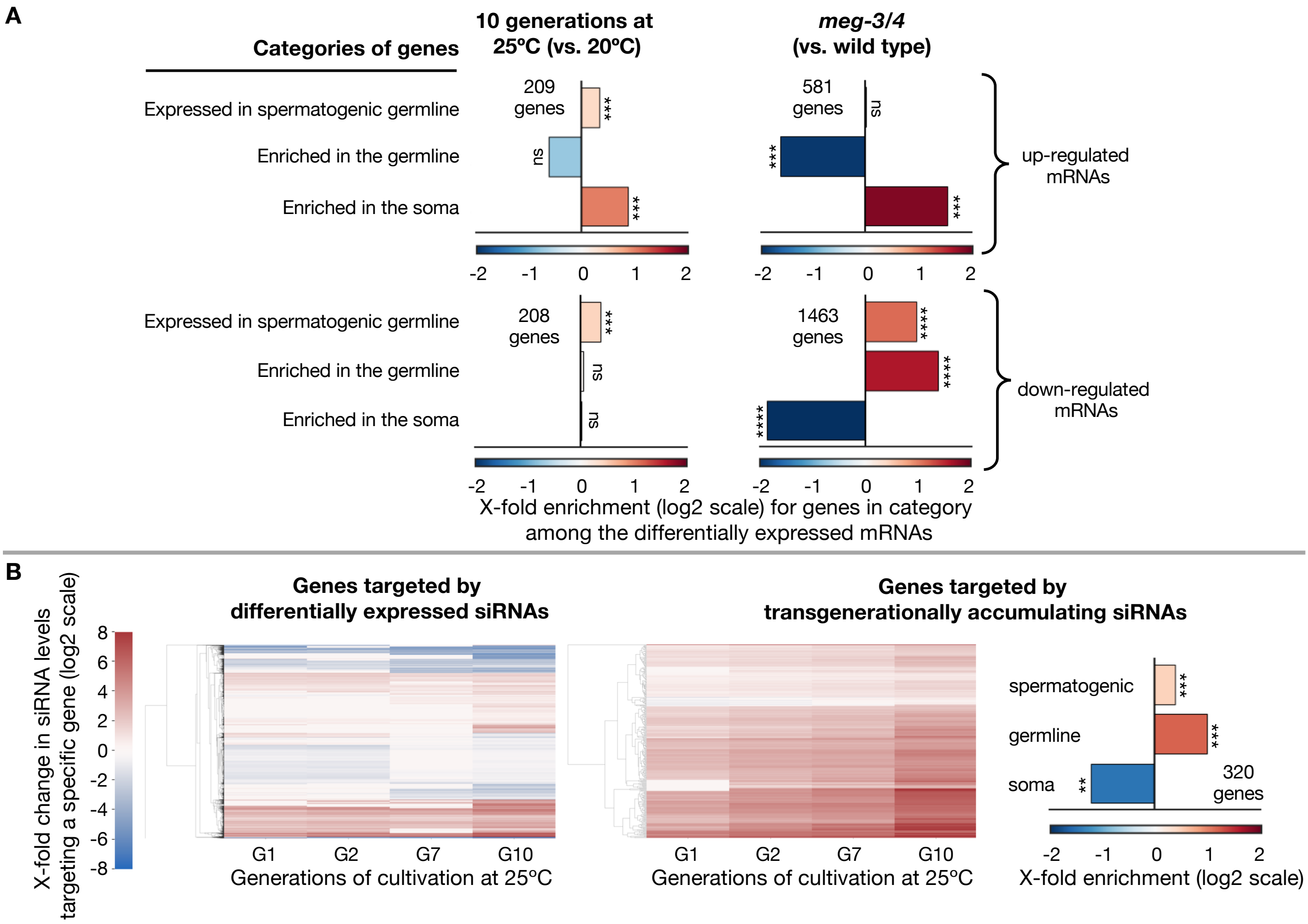


**Fig. S9. Additional analyses of siRNA and mRNA expression in prematurely attractive worms. (A)** Enrichment analysis of differentially expressed mRNAs. Enrichment analysis of mRNAs up-regulated (high panel) and down-regulated (low panel) in worms that were cultivated at 25ºC(*4*), and in *meg-3/4* mutants(*5*). Differentially expressed genes were analysed with Deseq2, adjusted *P* value < 0.1, and N of genes that passed the threshold appear next to the relevant panel. Shown are fold (log2 scale) enrichment results (observed/expected) in the lists of differentially expressed genes, for genes expressed in the spermatogenic germline(*6*), for genes enriched in the whole germline(*7*), and for genes enriched in the soma(*7*). Adjusted *P* values were obtained using a randomization test and corrected for multiple comparisons using the Benjamini–Hochberg step-up procedure. (**B**) Clustering of siRNA expression in worms grown at 25ºC for 1, 2, 7 & 10 generations. (Left) Hierarchical clustering analysis of 5223 genes targeted by differentially expressed siRNAs in worms grown at 25ºC for 10 generations compared to control. (Middle) A subset of 320 protein-coding genes targeted by transgenerationally accumulating siRNAs at 25ºC. siRNAs were considered "transgenerationally accumulating" if they displayed significantly higher differential expression levels in each time point compared to the previous one, and were significantly upregulated at the 10$^{th}$ generation. Color bar indicates the log2 fold changes in siRNA expression (compared to 20ºC grown worms, adj. $P < 0.1$). Non-significant fold-change values (adj. $P > 0.1$) appear in white. Data was generated by re-analysis of the available dataset by(*4*). (Right) Enrichment analysis for the subset of 320 protein-coding genes described above. ****$P < 10^{-4}$, ***$P < 0.001$, **$P < 0.01$, ns $P > 0.05$.

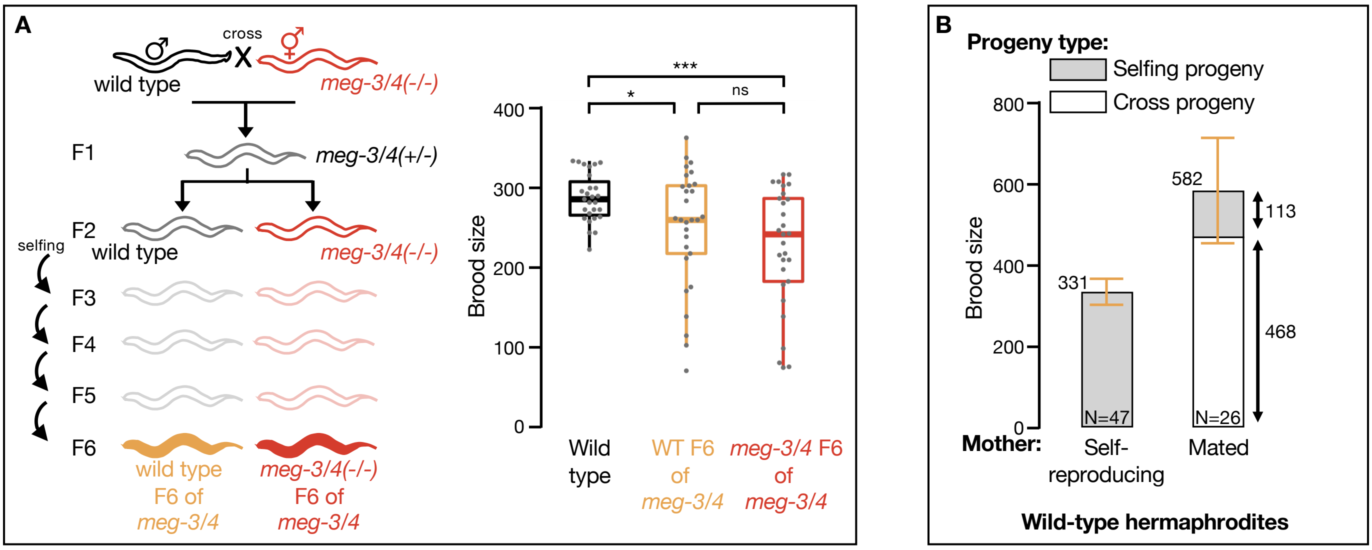


**Fig. S10. Brood size analyses.** (**A**) Brood size quantifications of wild-type (black), *meg-3/4* (red) and F6 wild-type descendants of *meg-3/4* hermaphrodites (yellow). As depicted in scheme, *meg-3/4* hermaphrodites were outcrossed with wild-type males, and homozygote F2s were isolated from the cross progeny. F6 descendants from both lineages were used for brood size experiments side by side with wild-type controls. All worms involved contained an integrated single-copy *mex-5p::gfp* transgene in their genetic background. Data collected over 3 independent experiments. Dots represent values for individual hermaphrodites. Boxplot (Tukey's style): median & IQR, whiskers extend to the most extreme value within 1.5xIQR from the 25$^{th}$ or 75$^{th}$ percentile. Welch's ANOVA, Games-Howell post-hoc correction for multiple comparisons. ***$P < 0.001$, *$P < 0.05$, ns $P > 0.05$. (**B**) Brood size quantification of self-reproducing or outcrossed wild-type hermaphrodites, used to generate the theoretical population genetics model (see Methods). Males used for outcrossing contained in their genome an integrated *myo-2p::gfp* transgene (pharyngeal muscles). GFP expression was used to determine mating status of the hermaphrodite and the genetic status of the progeny (selfing-progeny vs. outcrossed progeny). Data collected over 2 independent experiments. Bars: mean ± sd. Rounded averages for total brood size appear near the top left of bars, rounded averages for subgroups appear on the right. Genetic status of progeny is color coded.

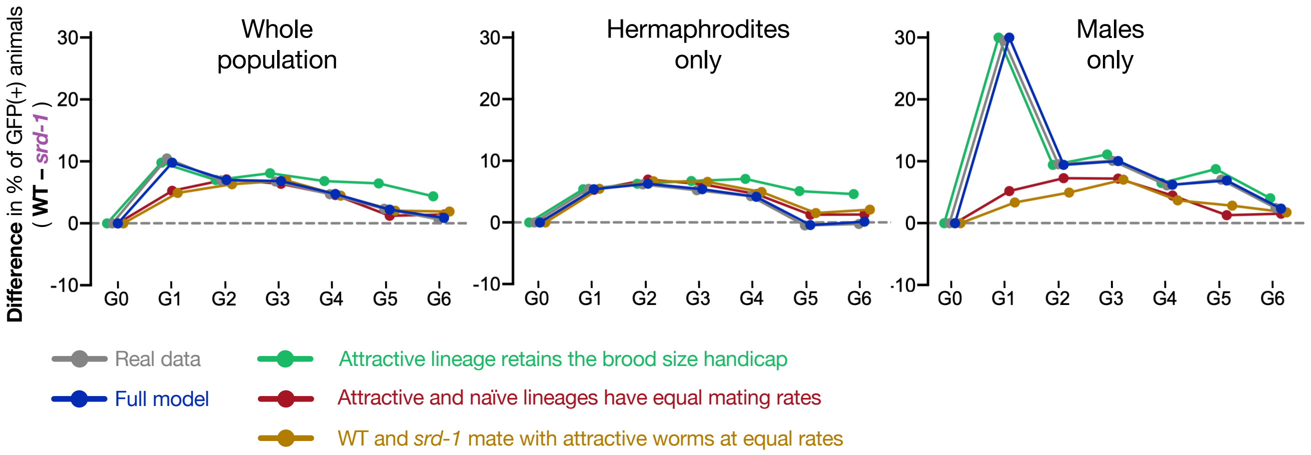


**Fig. S11. Computer simulations of the multigenerational competition experiment.** Computer simulations of the multigenerational competition experiment in which different scenarios were tested by relaxing the underlying assumptions of the experiment. Part of the data appears also in Figure 5C and is depicted here for convenience of visualization. On the y-axis is the difference in the proportion of *gfp(+)* worms (attractive lineage) in the population between the wild-type and *srd-1(-)* experiments. The experimental data from the multigenerational experiment (depicted in Fig. 5B) appear here in grey. The results of the full model (dark blue) reproduce the actual results (grey) with high accuracy. The main null hypothesis assuming equal mating rates for wild type and *srd-1* mutants is depicted in yellow. In this scenario the difference in *gfp(+)*% among males between the two experiments is expected to be substantially lower than the observed experimental results (see right panel). This is also true under the assumption that worms from the attractive lineage (wild-type descendants of *meg-3/4* worms) and the control naïve lineage have the same mating rates (red). We note that in order to accurately recapitulate the experimental results, we had to assume that the epigenetically inherited fitness handicap of the attractive lineage (see Fig. S9A) is weakened over generations. This assumption is consistent with the transgenerational recovery from RNAi defects witnessed in this lineage(*41*, *45*). In contrast, when assuming a constant fitness for the attractive lineage (green), a fairly constant difference between the lineages is observed throughout the generations (middle panel). This deviates from the experimental results where the difference in fitness between the two lineages is substantially reduced by the fifth and six generation.

#### References (Supplemental Information)


1. K. Nishimura, T. Fukagawa, H. Takisawa, T. Kakimoto, M. Kanemaki, An auxin-based degron system for the rapid depletion of proteins in nonplant cells. *Nat. Methods*. **6**, 917–922 (2009).
2. L. Zhang, J. D. Ward, Z. Cheng, A. F. Dernburg, The auxin-inducible degradation (AID) system enables versatile conditional protein depletion in C. elegans. *Dev.* **142**, 4374–4384 (2015).
3. B. A. Buckley, K. B. Burkhart, S. G. Gu, G. Spracklin, A. Kershner, H. Fritz, J. Kimble, A. Fire, S. Kennedy, A nuclear Argonaute promotes multigenerational epigenetic inheritance and germline immortality. *Nature*. **489**, 447–51 (2012).
4. K. I. Manage, A. K. Rogers, D. C. Wallis, C. J. Uebel, D. C. Anderson, D. A. H. Nguyen, K. Arca, K. C. Brown, R. J. C. Rodrigues, B. F. M. de Albuquerque, R. F. Ketting, T. A. Montgomery, C. M. Phillips, A tudor domain protein, SIMR-1, promotes sirna production at pirna-targeted mrnas in C. Elegans. *Elife*. **9** (2020), doi:10.7554/eLife.56731.
5. J. P. T. Ouyang, A. Folkmann, L. Bernard, C. Y. Lee, U. Seroussi, A. G. Charlesworth, J. M. Claycomb, G. Seydoux, P Granules Protect RNA Interference Genes from Silencing by piRNAs. *Dev. Cell*. **50**, 716-728.e6 (2019).
6. M. A. Ortiz, D. Noble, E. P. Sorokin, J. Kimble, A new dataset of spermatogenic vs. oogenic transcriptomes in the nematode Caenorhabditis elegans. *G3 Genes, Genomes, Genet.* **4**, 1765–1772 (2014).
7. J. Serizay, Y. Dong, J. Janes, M. Chesney, C. Cerrato, J. Ahringer, *Genome Res.*, in press, doi:10.1101/gr.265934.120.
8. I. Lev, I. A. Toker, Y. Mor, A. Nitzan, G. Weintraub, O. Antonova, O. Bhonkar, I. Ben Shushan, U. Seroussi, J. M. Claycomb, S. Anava, H. Gingold, R. Zaidel-Bar, O. Rechavi, Germ Granules Govern Small RNA Inheritance. *Curr. Biol.* **29**, 2880-2891.e4 (2019).
9. A. E. Dodson, S. Kennedy, Germ Granules Coordinate RNA-Based Epigenetic Inheritance Pathways. *Dev. Cell*. **50**, 704-715.e4 (2019).
